# Evidence for increased stress resistance due to polyploidy from synthetic autotetraploid *Caenorhabditis elegans*

**DOI:** 10.1101/2023.06.28.546823

**Authors:** Laetitia Chauve, Emma Bazzani, Clément Verdier, Liam Butler, Martha E. Atimise, Aoibhín McGarry, Aoife McLysaght

**Author notes:** equal second author contribution (Emma Bazzani, Clément Verdier).

## Abstract

Whole genome duplication (WGD) is a well-studied yet enigmatic phenomenon. While it has long been recognised as contributing numerous genes to many eukaryotic lineages and often implicated in evolutionary radiations, how these lineages overcome the known burdens of polyploidy is poorly understood. Circumstantial evidence of many WGD events coinciding with periods of otherwise mass extinction is consistent with the hypothesis that polyploidy is conditionally advantageous under stress conditions. While support for this comes from both theoretical work and field studies, direct evidence is lacking, especially in animals. Here we compare diploid and neo-tetraploid *Caenorhabditis elegans* and show that tetraploid animals exhibit increased resilience under specific stress conditions related to temperature changes. Most notably, under severe cold stress gravid neo-tetraploids massively escape cold-induced death, and generate more progeny, of similar quality, than diploid animals. This is the first demonstration of the effects of polyploidy on stress resistance and physiology in animals.

---

Whole Genome Duplication (WGD), or polyploidisation, is an unusual mutational event where the entire genome becomes repeated within the nucleus of the cell. WGD can arise either from hybridisation between genomes of two related species (allopolyploidisation) or from doubling the entire set of chromosomes of a given species (autopolyploidisation), perhaps due to an error during meiosis. WGD is a major evolutionary force that has had profound and lasting effects on the genomes of animals, plants and protists^1^.

Polyploidy is a relatively common mutational event, including in healthy and diseased somatic cells (endopolyploidy)^2–7^, but rare over evolutionary times. Genome duplication has been identified in most eukaryotic lineages^8^. WGD is common in plants, where it has been linked to speciation events, and the ability to withstand periods of stress^9,10^. It is rarer in animals, with an estimated frequency of less than 1%^11,12^ but this low number has a potentially high impact, most notably around the establishment of the vertebrate lineage^13–16^ and at the base of teleost fish^17^. The success of these ancestrally polyploid (‘paleopolyploid’) lineages is self-evident from their mere existence. However, this throws open the important question of how and why these lineages survived through what is often referred to as the evolutionary ‘dead end’ of polyploidy. While in allopolyploids it may be advantageous to bring together the best of two different genomes, a phenomenon known as hybrid vigour or heterosis, the short-term benefits of autopolyploidy are harder to discern. Nevertheless, it is estimated that about half of the WGD events in angiosperms originate from autopolyploidisation^18^, and at least some of the ancestral WGD events in vertebrates are inferred to have been autopolyploidies^19,20^.

The WGD events that have been well-studied are generally very ancient and little is known about the short-term or immediate consequences of polyploidisation on the genome or physiology. Studies in plants show that autopolyploidisation poses significant challenges, particularly during meiosis, where issues such as chromosome segregation errors and genomic instability arise^1,21,22^. Neopolyploid organisms often display reduced fertility, which is commonly attributed to aneuploidy, although fertility defects in nascent tetraploids can also stem from morphological abnormalities^23^. In line with this, genomic analyses of a few recently established plant autopolyploids, highlight signatures of adaptation to meiosis and DNA damage^24,25^.

However, what factors contribute to the success of a polyploid lineage remains unknown, particularly in animals. One attractive though debated hypothesis states that polyploidy might increase tolerance to stressful environments and be adaptive in the short-term^9^. It is well established that stress itself (*e.g.,* heat or cold) can lead to autopolyploidy via the generation of unreduced (*i.e.*, diploid) gametes by failure of germline cell division after meiotic replication^11^. Therefore, a legitimate question is whether autopolyploidy might have an adaptive value under stressful conditions or whether it is merely a consequence of stress.

In plants, WGD events are often associated with periods of climatic change and environmental instability^1,9^. Additionally, experimental evolution studies reveal that populations of autotetraploid yeast fix beneficial mutations faster than diploids^26^. Although initially more sensitive to elevated temperatures, lab-evolved tetraploid yeast become more resilient to heat stress^27^. Modelling approaches revealed increased variation of gene regulatory networks following polyploidy, which may prove advantageous under environmental turmoil^28^. These studies point to a potential adaptive benefit, which has also been suggested for amphibians and fish^29^. Indeed, the distribution of the Australian autotetraploid burrowing frog *Neobatrachu*s suggests that autotetraploids are better adapted to harsher environments^1^ perhaps through facilitating gene flow^30^.

However, to unlock these longer-term benefits nascent polyploids must first overcome the challenges associated with polyploidy. Could it be that WGD provides immediate benefits under specific conditions? In plants, autopolyploidisation is associated with increased abiotic stress resistance^31^ and resistance to pathogens^32,33^ and increasing ploidy in yeast can be advantageous under specific stress conditions^34^. However, the immediate effects of WGD in animals remain currently unexplored.

To investigate whether autopolyploidy has an adaptive value in the short term in animals, we sought to explore the consequences of induced autopolyploidy on physiology and stress responses in the nematode *Caenorhabditis elegans,* where it is possible to generate neo-autotetraploids by transiently knocking down the cohesin complex component *rec-8* by RNA interference for two generations^35^. *C. elegans* provides several advantages as a model to study the consequences of polyploidy^2^: derived tetraploid animal lines are stable (producing mostly tetraploid offspring) and fertile enough to work with; and with its mostly self-fertilizing hermaphrodite mode of reproduction, it provides isogenic genetic backgrounds for comparison, differing only by ploidy levels.

In animals, the impact of unscheduled autotetraploidy on physiology, particularly in the context of stress, remains unexplored. Here we generated autotetraploid animals and exposed these synthetic tetraploids to various stresses. We show that, as expected, unscheduled autotetraploidy is deleterious under regular growth conditions: neotetraploids have a shorter lifespan and decreased fertility.

Exposing neotetraploid *C. elegans* to stressful environments revealed altered phenotypes under specific stress conditions related to temperature changes. While neotetraploids exhibited similar survival on pathogenic bacteria, their lifespan becomes similar to WT when the temperature increases. Neotetraploids displayed a modest increase in heat stress resistance, accompanied by altered nuclear localisation of the DNA locus of the highly inducible molecular chaperone *hsp-16.2*. This did not, however, translate into an altered induction pattern of highly inducible molecular chaperones. Remarkably, when exposed to severe cold stress followed by recovery, gravid neotetraploid animals massively escaped cold-induced death. Additionally, following recovery from cold stress, neotetraploids produced more progeny of equal quality compared to their diploid counterparts. This suggests a potential adaptive value of autotetraploidy under cold stress conditions. Lastly, we checked whether known mechanisms associated with cold-induced death in gravid diploids were altered in tetraploids. Our data points to novel, yet unknown, mechanisms underlying escape from cold-induced death in tetraploids.

## RESULTS

### Fitness of tetraploids decreased under regular conditions

To assay the physiological consequences of unscheduled autopolyploidy in *C. elegans*, we generated neotetraploid animals using *rec-8* RNA interference^35^. We obtained stable lines engendering almost exclusively tetraploid offspring, with only rare events of reversion to diploidy. Tetraploids were selected in the F2 progeny based on body-size increase^35,36^ (**Figure 1C**). We confirmed polyploid status by checking for doubling of chromosome bivalents in late-stage oocytes at the diakinesis stage (**Figure 1A-B**), similarly to previous work^35^.

**Figure 1.**
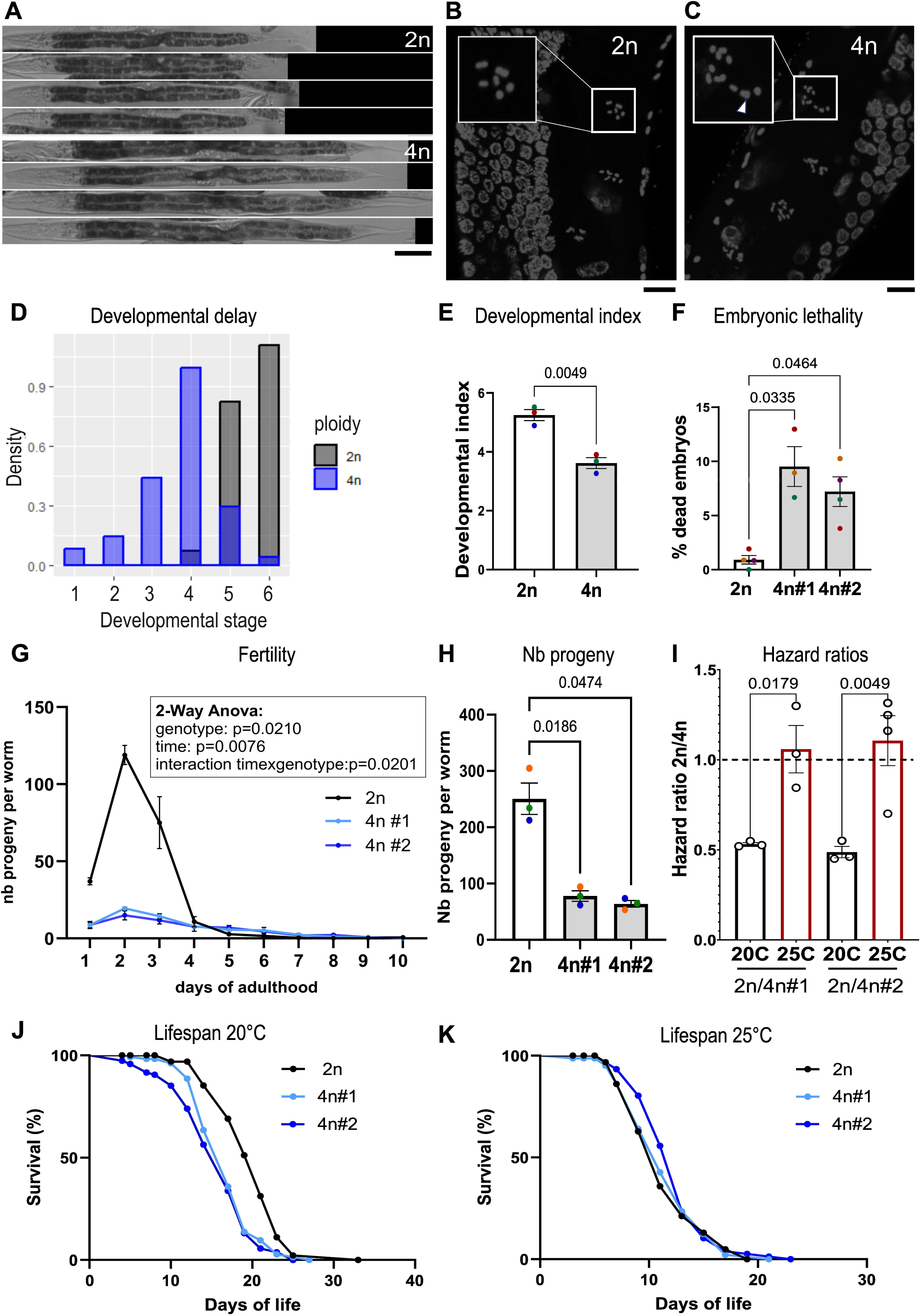
Synthetic autotetraploid *C. elegans* animals exhibit decreased fitness under regular conditions. **(A)** Images of diploid (2n, top) N2 and tetraploid (4n, bottom) MCL2 animals at L4 stage, showing length differences. Scale bar is 100 µm. **(B-C)** Micrographs of germlines of diploid N2 (B) and derived MCL2 tetraploid animals (C) stained with DAPI. The most mature unfertilized oocyte nuclei before the first meiotic division are highlighted with a box and show 6 bivalent chromosomes in 2n (B), and 12 bivalents in 4n (C). Arrow in C indicates two partially overlapping chromosomes bivalents. Maximum intensity projection, Confocal, objective 63X. Scale bar is 10µm. **(D)** MCL2 (4n) tetraploid animals are developmentally delayed. Developmental stage at 65h post-egg-laying synchronisation at 20°C. Stages: L1, L2, L3, L4, young adult (YA), and gravid adult (GA) are numbered from 1 to 6 respectively. **(E)** Tetraploid MCL2 (4n) are delayed compared to diploid animals. A developmental index was calculated by multiplying frequencies of worms at a particular stage (at 65h post synchronization at 20°C), by numbered developmental stages (i.e. L1=1). Paired t-test: p-value=0.0049 (*). **(F)** Derived tetraploids MCL1 (4n#1) and MCL2 (4n#2) exhibit higher rates of embryonic lethality at 20°C than diploid WT N2 (2n). Mixed effect analysis with Geisser-Greenhouse correction and Dunnet’s multiple comparison test; p-value=0.0235 (*). **G)** Derived tetraploids MCL1 (4n#1) and MCL2 (4n#2) exhibit decreased fertility compared to WT N2 (2n) animals. **(H)** Average number of progeny per worm: N2: 231.5, MCL1: 61.46, MCL2: 65.97. Two-way ANOVA analysis. **(I)** Hazard ratios of 2n/4n raised at either 20°C or 25°C growth temperature for derived tetraploids MCL1 (4n#1) and MCL2 (4n#2). At 20°C, the probability of death is two times lower for 2n than for 4n, whereas at 25°C the hazard ratios are similar (∼1) for 2n/4n. See also table S3. **(J)** Lifespan at 20°C of WT N2 (2n) and derived tetraploid lines MCL1 (4n#1) and MCL2 (4n#2). Number of animals assayed: 148-150 per genotype. Log-rank (Mantel-Cox) test: N2 (2n) vs MCL1 (4n #1): p-value<0.0001 (****); N2 (2n) vs MCL2 (4n #2): p-value<0.0001 (****). **(K)** Lifespan at 25°C of WT N2 (2n) and derived tetraploid lines MCL1 (4n#1) and MCL2 (4n#2). Number of animals assayed: 148-150 per genotype. Log-rank (Mantel-Cox) test: N2 (2n) vs MCL1 (4n #1): p-value=0.8008 (ns); N2 (2n) vs MCL2 (4n #2): p-value=0.0869 (ns). In all panels, colours indicate matching independent biological replicates. Error bars = SEM.

Phenotypic characterization under regular growth conditions (20°C) showed that neoautotetraploid animals are developmentally delayed (**Figure 1D-E**) exhibiting more stage heterogeneity at 65h post egg-laying synchronization than diploids. The delay in reaching adulthood for most tetraploid animals is estimated to be between 8-10h at 20°C. Autopolyploidisation severely affected the number of offspring, which reduced to 28.5% of WT progeny numbers (**Figure 1G-H**). As polyploidisation can have deleterious consequences for chromosome segregation^1,21,22^, we monitored embryonic lethality, often caused by aneuploidies (**Figure 1F**). Our analysis showed that neotetraploids exhibit a higher rate of embryonic lethality (between 7-9%) than diploids (around 1%). However, this increased rate of embryonic lethality seems insufficient to explain the 70% decrease in fertility. Additionally, lifespan was significantly decreased in two independent lines of neoautotetraploids at 20°C (**Figure 1I-K**). Altogether, these results show that, under regular growth conditions, unscheduled autopolyploidisation has deleterious consequences on fitness and negatively affects development, fertility, and lifespan.

### Tetraploids exhibit similar resistance to pathogenic bacteria

In plants, autopolyploidy confers resistance to pathogenic bacteria by constitutively activating plant defences^32,37^, and modelling approaches predict better resistance of polyploids to pathogens and parasites^38^. By contrast, synthetic triploid rainbow trout and Atlantic salmon are more susceptible to viruses, bacteria and parasites^39^. We tested the resistance of synthetic autotetraploid *C. elegans* to its most studied pathogen, *Pseudomonas aeruginosa*. *P. aeruginosa* is an opportunistic pathogen of animals, insects, nematodes, and plants. *P. aeruginosa* kills *C. elegans* in several ways, depending on the environment^40,41^. Under conditions of low osmolarity, *P. aeruginosa* colonises the intestine and kills nematodes within a few days (slow-killing assay).

We monitored resistance to pathogenic bacteria in animals of different ploidies using *P. aeruginosa* strain PAO1. As exposure to *P. aeruginosa* leads to a matricide “bagging” phenotype in gravid adults, we performed the experiments in sterile nematodes, either upon exposure to FUdR (5-Fluoro-2’-deoxyuridine) which inhibits cell divisions in the progeny (**Figure 2A**), or using the gonadless temperature sensitive genetic background *gon-2(q388);gem-1(bc364)*^42^ (**Figure 2B**). We observed similar resistance of neotetraploids compared to diploids under both conditions (**Figure 2A-B**), suggesting that autotetraploidy does not alter the response to pathogenic *P. aeruginosa* in *C. elegans*.

**Figure 2.**
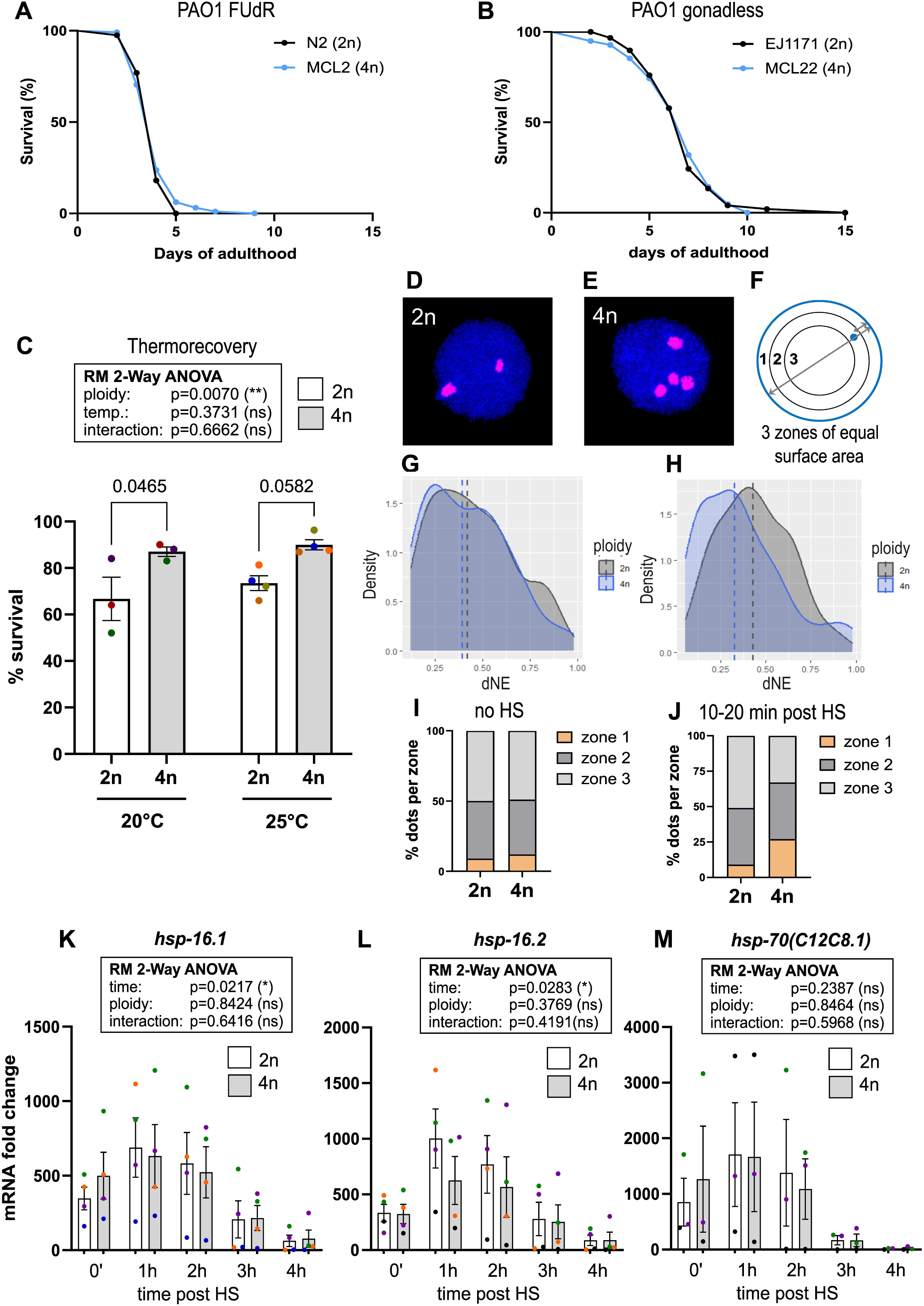
Synthetic autotetraploid *C. elegans* display similar resistance to pathogenic bacteria and improved resistance to heat stress. **(A)** Survival of WT diploid N2 (2n) and derived tetraploid MCL2 (4n #2) animals raised on pathogenic *Pseudomonas aeruginosa* PAO1 bacteria from L4 stage. Animals were grown on plates containing DNA synthesis inhibitor FUdR to prevent egg-hatching and bagging, common upon PAO1 exposure. Experiment was performed at 25°C. p-value N2 (2n) vs MCL2 (4n) = 0.7546 (ns). **(B)** Survival of gonadless WT diploid EJ1171 (2n) *gon-2(q388); gem-1(bc364)* temperature-sensitive (ts) mutants and derived gonadless tetraploid MCL22 (4n) animals raised on pathogenic *Pseudomonas aeruginosa* PAO1 bacteria from L4 stage on. Experiment performed at 25°C. p-value EJ1171 (2n) vs MCL22 (4n) = 0.9263 (ns). **(C)** Survival upon thermorecovery of N2 (2n) and derived tetraploid MCL2 (4n) animals exposed to 4h heat stress at 35°C, followed by 20h recovery at their respective growth temperatures. Independent biological replicates are coloured. **(D-E)** Micrographs of early embryonic nuclei (∼50 cells stage) from GW615 (2n) or derived tetraploid MCL7 (4n) animals carrying both transgenes *baf-1*p::GFP-LacI and *hsp-16.2p-*LacO^44^. Nuclei were imaged at 63X obj. on a confocal and visualised using Imaris software, with nuclear fluorescence from *baf-1p::*GFP-LacI in blue and *hsp-16.2*p-LacO DNA loci in pink. **(F)** Schematic representation of *hsp-16.2p*-LacO DNA locus intranuclear position measurements according to the method developed in^44,49^. For each dot, the distance to the nuclear envelope (dNE) was measured as well as the diameter of the nuclear plane. **(G-H)** Density plots representing the distance to the nuclear envelope (dNE) of *hsp-16.2p*-LacO loci in the absence of heat shock (HS) (G) or 10-20 minutes following a 10 minute HS at 34°C (H) in GW615 (2n) or derived tetraploid MCL7 (4n). Data in (G): GW615: 49 nuclei (96 dots), MCL7: 20 nuclei (54 dots). Unpaired t-test GW615 vs MCL7: p-value=0.5715. Data in (H): GW615: 47 nuclei (88 dots), MCL7: 38 nuclei (119 dots). Unpaired t-test GW615 vs MCL7: p-value= 0.0204. Dashed lines in G and H represent the median. **(I-J)** Classification of *hsp-16.2p*-LacO genomic DNA loci positions in 3 zones of equal surfaces within the nuclear plane, as defined in^44^ in the absence of HS (I), or following a short HS of 10 minutes at 34°C (J), for diploid GW615 and tetraploid MCL7 nuclei. (I): Chi-square GW615 vs MCL7: p-value = 0.8638 (ns). (J): Chi-square GW615 vs MCL7: p-value=0.0019 (**). **(K-M)** mRNA expression levels following a 30-minute heat shock at 34°C (HS) of *hsp-16.1* (K), *hsp-16.2* (L) and *hsp-70(C12C8.1)* (M). RM two-way ANOVA with Geisser-Greenhouse correction and Šídák’s multiple comparison test. For each ploidy, the mRNA levels were normalized such that levels without HS=1. Fold change mRNA expression levels of mRNA targets were normalized to 5 housekeeping genes (Y45F10D.4, *pmp-3, lap-2, klp-12, and act-1*). The p-values for each factor of the two-way ANOVA are indicated on the graph. In all panels, colours indicate matching independent biological replicates. Error bars = SEM.

### Tetraploids are slightly more thermotolerant than diploids

While unscheduled autotetraploidy negatively affects lifespan under regular growth conditions at 20°C (**Figure 1I, K**), there was no deleterious effect on lifespan under conditions of mild heat stress at 25°C (**Figure 1J-K**). The improvement in the lifespan of neotetraploids with increasing temperature prompted us to assess the heat stress response in neotetraploids. The heat stress response is a highly conserved response relying on the transcriptional activation of molecular chaperones (also called *heat shock proteins – hsps)* upon heat stress, and inhibition of general translation to maintain and restore cellular protein homeostasis^43^. After recovery from severe heat stress (thermorecovery) neotetraploid animals raised at either 20°C or 25°C exhibited a modest increase in survival (p-value ploidy effect =0.0070, **Figure 2C**), suggesting that neotetraploids can mount a stronger heat stress response.

Molecular chaperones are comprised of several gene families, some of which are constitutively expressed (i.e. *hsp-1, hsp-90*) while others are expressed at low levels under normal conditions and highly induced and attenuated following proteotoxic stress, such as *hsp-16* (small *hsp* family) and *hsp-70* (C12C8.1). Molecular chaperones are essential to maintain protein homeostasis, *i.e.,* the proper balance between protein synthesis, folding, trafficking, and degradation, collectively referred to as proteostasis. Our data revealed similar expression levels of constitutive molecular chaperones in animals of different ploidies under regular conditions (**Figure S2D**). This indicates that tetraploid worms do not show signs of proteotoxic stress under regular conditions. To further investigate proteostasis in tetraploids, we used characterised proteostasis sensors: the temperature-sensitive *unc-54* mutation which leads to paralysis when proteostasis is compromised in muscles; and multicopy proteostasis sensors polyQ35 and polyQ44. Our results using the *unc-54* mutation at either 15°C (permissive) or 25°C (restrictive temperature) revealed overall no significant difference in motility between diploids and tetraploids, suggesting similar proteostasis capacity (**Figure S2J, K**). However, similar experiments using multi-copy transgenic proteostasis sensors show contrasting results, with tetraploids exhibiting more protein aggregation in both muscles and intestine (**Figure S2 A-C, S2 F-I**). These discrepancies raise the possibility that the regulation of multicopy transgenes (such as *unc-54p::Q35::YFP* and *vha-6p::Q44::YFP*) could be altered by tetraploidy. Indeed, we observed differences in the expression levels of the *hsp-90p*::GFP single-copy/multicopy transgene in animals of different ploidies (**Figure S3**).

We wondered whether and how autotetraploidisation could affect the heat stress response, as suggested by increased survival from thermorecovery The DNA locus of highly inducible molecular chaperones, such as *hsp-16.2* is located in proximity to the nuclear envelope and the nuclear pore, for rapid mRNA export upon heat shock^44^. The effects of increased cell size due to polyploidy on sub-cellular localisation and dynamics is currently unknown. Polyploidy is presumed to alter the ratio of the nuclear envelope surface to nuclear volume: assuming the nucleus to be a sphere, when the volume of the nucleus is doubled the nuclear envelope is only increased by 1.6 fold^45^ (though observations in animals and plants suggest that this is a non-trivial relationship^46–48^).

We hypothesised that the altered heat shock response in neotetraploids could be due to altered positioning of the *hsp-16.2* DNA locus relative to the nuclear envelope. To explore this, we generated neo-autotetraploid animals carrying the in-vivo tracking system for the *hsp-16.2* DNA locus: *hsp-16.2p*::LacO/*baf-1p*::GFP-LacI^44^. As depicted in **Figure 2D-E**, we measured the nuclear positioning of *hsp-16.2* loci relative to the nuclear envelope in early embryonic (∼50 cell stage) nuclei by classifying the position of each *hsp-16.2* locus into three zones of equal surface within the nuclear plane, (**Figure 2F**) as per ref^49^. This analysis showed that tetraploidy does not affect *hsp-16.2* DNA locus positioning relative to the nuclear envelope in the absence of heat shock (**Figure 2G, I)**. However, following a short pulse of heat shock (10 minutes at 34°C), *hsp-16.2* loci were more frequent in zone 1 and more closely associated with the nuclear envelope than in diploids (**Figure 2H, J**). To determine whether altered nuclear localisation of *hsp-16.2* following heat shock in tetraploids could affect its expression levels, we checked its induction kinetics by quantitative Real-Time PCR (qRT-PCR). We monitored the kinetics of induction following a short heat shock pulse (30 min at 34°C) for the highly inducible molecular chaperones *hsp-16.1* and *hsp-16.2,* and *hsp-70(C12C8.1).* Our analysis revealed overall similar kinetics and amplitude in the transcriptional activation and attenuation of *hsp-16.1* and *hsp-16.2 and hsp-70* upon heat shock (**Figure 2K, M**).

Altogether, these data reveal that, while proteostasis capacity seems similar in animals of different ploidies under regular conditions, neotetraploids exhibit a modest increase in thermotolerance and that *hsp-16.2* is more closely associated with the nuclear envelope upon heat shock without any detectable consequences on highly inducible *hsps* induction kinetics.

### Gravid adult tetraploids escape cold-induced death under severe cold stress

Autopolyploidisation is associated with increased tolerance to cold in plants, both in established polyploids^50^, or in neoautopolyploids^51,52^, whereas the response to cold stress remains poorly understood in polyploid animals.

We assayed resistance to severe cold stress in neotetraploid animals. Plates containing worms were placed in a box of ice for four hours, followed by 20 hours of recovery at 20°C (**Figure 3A**). While ploidy did not significantly affect survival after cold recovery at the L4 stage (**Figure 3B**), there was a dramatic difference in gravid adults at day 2 of adulthood. Diploid day 2 adults massively died (∼5% survival) while 80-90% of tetraploid day 2 adults survived (**Figure 3B**). This difference in survival was still observable two days later, at 72h post cold shock (**Figure S5B**). Diploid revertants, derived from neotetraploids, phenocopied diploid WT day 2 adults, suggesting survival from cold recovery is associated with tetraploid state. At the time of scoring, the matricide “bagging” phenotype was observed in day 2 adult diploids. To ensure death of diploids was caused by exposure to cold rather than by internal hatching, we performed the cold recovery assay in day 2 adults using FUdR, thus preventing meiosis. We could still detect a significant difference in survival (**Figure 3B**). Cold-induced death at day 2 of adulthood depends on the presence of a fully functional gonad, as temperature sensitive gonadless diploid *gon-2(q388);gem-1(bc364)* mutants survived cold recovery as day 2 adults at a similar rate to neotetraploids (**Figure 3B**).

**Figure 3.**
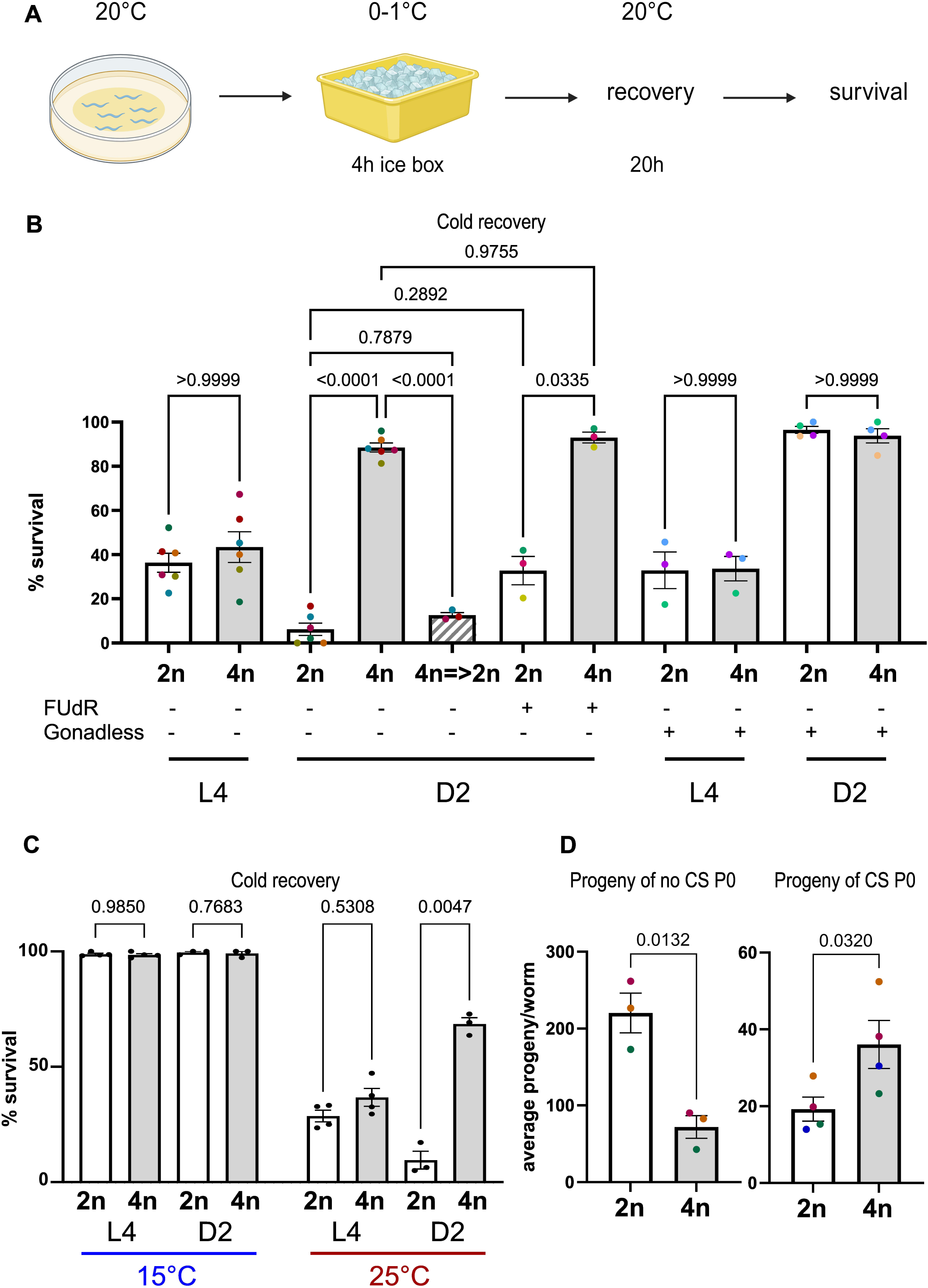
Synthetic autotetraploid *C. elegans* escape cold-induced death at the adult stage and produce more progeny than diploids. **(A)** Schematic overview of cold recovery assay. Plates containing animals raised at 20°C were placed in ice for 4h, followed by a 20h recovery period at 20°C. **(B)** Survival upon cold recovery of diploid N2 (2n), derived tetraploid MCL2, diploid revertants from neotetraploids (4n=>2n) at L4 or day 2 adult stage (D2). N2 (2n) and MCL2 were also grown on plates containing DNA synthesis inhibitor FUdR to prevent egg-hatching and bagging. The last four columns on the right indicate survival upon cold recovery of gonadless diploid EJ1171 (2n) *gon-2(q388); gem-1(bc364)* temperature-sensitive mutants and derived gonadless tetraploid MCL22 (4n) at L4 or Day 2 adult stage (D2). Brown-Forsythe and Welch’s ANOVA test with Dunnet’s T3 multiple comparison test with individual variances computed for each comparison. P-value Brown-Forsythe test: <0.0001(****), p-value Welch’s test: <0.0001 (****). **(C)** Survival upon cold recovery of diploid N2 (2n) and derived tetraploid MCL2 (4n) at L4 or Day 2 adult stage. Animals were either raised at 15°C (cold acclimatation) or at 25°C. Mixed-effects model analysis with Greenhouse-Geisser correction and uncorrected Fisher’s LSD was run separately at each growth temperature. At 15°C, p-value ploidy effect=0.5531 (ns), p-value stage effect=0.3399 (ns), and p-value interaction= 0.9242 (ns). At 25°C, p-value ploidy effect=0.0002 (***), p-value stage effect=0.1193 (ns) and p-value interaction= 0.0006 (***). **(D)** Average number of progeny per P0 day 2 adult worms of diploid N2 (2n) or derived tetraploid MCL2 (4n #2) under regular conditions (left), or exposed to cold recovery (right). Mixed-model analysis with Geisser-Greenhouse correction: p-value=0.0069(**). Adjusted p-value with Šídák’s multiple comparisons test are indicated on the graph. In all panels, colours indicate matching independent biological replicates.

Recovery from severe cold stress is affected by cold acclimation at lower growth temperature (15°C)^53^. Cold acclimation is controlled by neuron-intestine hormonal signalling from a subset of thermosensory head neurons^53,54^. Under growth at a lower temperature, neuroendocrine insulin and steroid hormone signalling from the ASJ neuron leads to the activation of insulin signalling in the intestine and lipid composition changes promoting cold resistance^54^. We assayed cold recovery at different growth temperatures. While growth at 25°C resulted in a similar phenotype as observed at 20°C, cold acclimation at 15°C promotes close to 100% survival after cold recovery in a similar manner in diploids and tetraploids, at both L4 and day 2 adult stage (**Figure 3C**).

### Cold-shocked tetraploids produce more progeny of similar quality compared to diploids

Fertility is strongly compromised in neotetraploids under normal conditions. As neotetraploid gravid adults escape cold-induced death, while most diploid adults die within 16h after cold shock, we wondered how the output number of progeny of cold-shocked neotetraploids would compare to the progeny of cold-shocked diploid animals upon recovery from severe cold stress. Our analysis revealed that the overall number of progeny of cold-shocked tetraploids is 2-fold higher than that of cold-shocked WT diploids, suggesting a potential adaptive advantage of neotetraploidy under cold stress conditions (**Figure 3D**). In diploids cold-shocked as day 2 adults, the average number of progeny is reduced by 12-fold compared to non-cold-shock day 2 adults (**Figure 3D**), with the progeny coming from eggs laid within 0 to ∼5h hours after the end of cold shock and from internal hatching when P0 diploids die around 10-12h post cold shock. On the other hand, in tetraploids, the average number of progeny following cold shock is only reduced by 2-fold, compared to non-cold shock conditions (**Figure 3D**).

Cold-induced death is proposed as a terminal investment response to favour progeny survival to the detriment of parents, with embryos of cold-shocked diploids better resisting subsequent cold shock. We sought to determine whether the progeny of cold shocked tetraploids, which escape cold-induced death, might exhibit decreased fitness under regular conditions compared to the progeny of cold shocked diploids. Upon recovery from cold shock, progeny were collected at different time points (**Figure 4A**). Embryonic lethality was severely increased in the F1 progeny of cold-shocked animals (**Figure 4B**), however, neotetraploid F1 animals seemed slightly less affected, however it was not significant (p-value ploidy = 0.1392).

**Figure 4.**
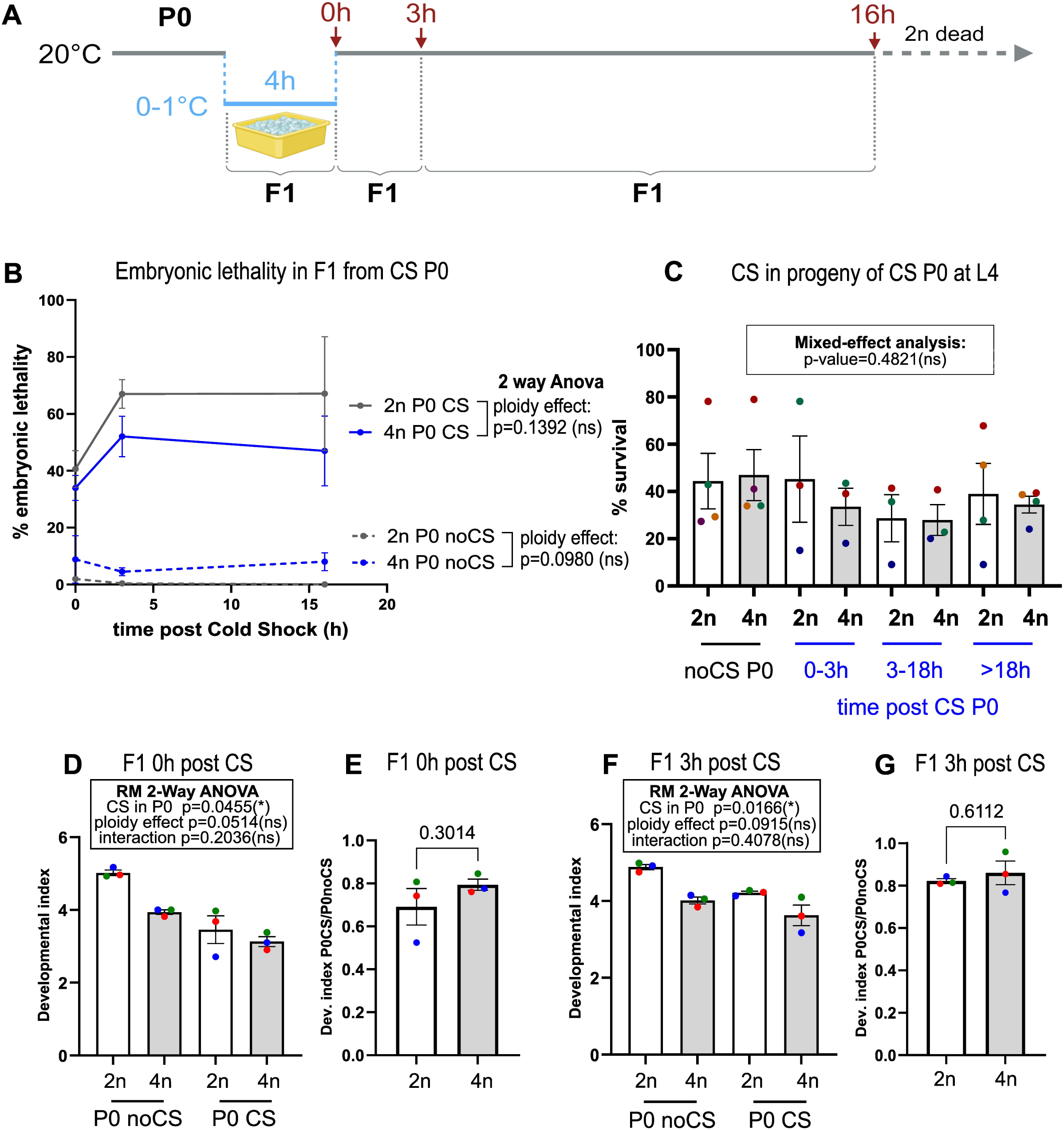
The progeny of cold-shocked autotetraploid animals is of similar quality than that of cold-shock diploid animals. **(A)** Schematic overview of different time points after cold shock (CS) and recovery at which F1 progeny of diploid and tetraploid cold shock P0 was assayed. Around 16h post cold shock, the majority of P0 diploid CS was dead. **(B)** Percentage of embryonic lethality in the progeny of diploid N2 (2n, grey) and derived tetraploid MCL2 (4n, blue) in the absence of CS (dashed lines) or following CS (full lines), at different time points after CS. RM Two-way ANOVA across all time points with Geisser-Greenhause correction and multiple correction test. Without CS: p-value ploidy effect: 0.0980 (ns), p-value time=0.6852 (ns), p-value ploidy x time: 0.7684 (ns). After CS: p-value ploidy effect: 0.1392 (ns), p-value time= 0.2652 (ns), p-value ploidy x time= 0.5605 (ns). **(C)** Survival upon cold recovery of F1 progeny of cold-shocked P0 diploid N2 (2n) or derived tetraploid MCL2 (4n) animals at the L4 stage. F1 progeny were assayed at different time points after CS of P0 animals. Mixed effect analysis: p-value N2 vs MCL2= 0.4489 (ns). Pairwise adjusted p-values (Šídák’s multiple comparisons test) are indicated on the graph. **(D-G)** Developmental index of the F1 progeny of CS P0 diploid N2 (2n) or derived tetraploid MCL2 (4n). (F1 progeny was collected right after CS of P0 (D-E) or at 3h post CS of P0s (F-G). As tetraploid MCL2 are developmentally delayed compared to diploid N2 animals, the developmental index of the F1 progeny cold-shocked of diploid and tetraploid was compared to the progeny of non-cold-shocked diploid and tetraploid animals respectively. (D-F). A developmental index was calculated by multiplying frequencies of worms at at a particular stage by numbered developmental stages at 65h post synchronisation at 20°C (i.e. L1=1). (D) Two-way ANOVA: p-value poidy effect=0.0191(*), p-value CS status in P0= 0.0067(**), p-value interaction ploidy x CS status in P0= 0.1114 (ns). Pairwise adjusted p-values from Šídák’s multiple comparison tests are indicated on the graph. (F) Two-way ANOVA: p-value poidy effect=0.0507(ns), p-value CS status in P0= 0.0741(ns), p-value interaction ploidy x CS status in P0= 0.0871(ns). Pairwise adjusted p-values from Šídák’s multiple comparison tests are indicated on the graph. (E-G) Ratio of developmental index of the progeny of cold-shocked P0s over the developmental index of the progeny of non-cold-shocked P0s. P-values from paired t-tests are indicated on the graphs in E and G. In all panels, colours indicate matching independent biological replicates. Error bars= SEM.

As another measure of fitness, we assayed developmental stage of synchronized F1 progeny from P0 at either 0h (zero hours) or 3h post cold shock (when P0 diploids were still alive). As tetraploids are developmentally delayed compared to their diploid counterparts (**Figure 1D-E**), we compared the stages of cold-shocked progeny to non-cold-shocked animals of the same ploidy. At 0h and 3h post-cold shock, the developmental index of both cold-shocked diploids and tetraploids was reduced (**Figure 4D, F**). The developmental delay in the F1 progeny of cold-shocked parents was not different in diploids and tetraploids (at 0h post CS: p-value = 0.3014 **Figure 4E**, at 3h post CS: p-value = 0.6112 **Figure 4G**). To test whether the progeny of cold-shocked P0 could exhibit increased recovery after CS, we then exposed the progeny of cold shocked P0 animals of different ploidies, collected at different time points, to a cold shock at the L4 stage. However, we could not detect any difference in survival of the progeny between the two ploidies (**Figure 4C**). Altogether, these data indicate that gravid neotetraploids escape cold-induced death and produce twice as much progeny than diploids following cold shock. The fitness of the progeny of neotetraploids is similar, if not slightly better, compared to the progeny of their diploid counterparts.

### The survival of CS tetraploid gravid adult is uncoupled from cold-induced intestine-to-germline lipid relocalisation

In diploids, recovery from severe cold stress leads to lipid relocalisation from the intestine to the embryos^55^, at the expense of parental survival. We investigated whether tetraploids exhibited differences in fat content and whether tetraploids recovering from cold shock were exempt from intestine-to-germline lipid movements, potentially explaining their survival from severe cold shock^55^. Under regular conditions, we found that fat stores were overall similar in gravid adults of different ploidies, as monitored using the fluorescent dye BODIPY 493/503 **(Figure 5A-B** and **Figure 5E)** or using Oil Red O (ORO staining **(Figure S4C-E)**. At 10h recovery from cold shock, our BODIPY analysis revealed an overall decrease in fat stores in both diploid and tetraploid animals, which was more pronounced in the intestine, while the embryos retained most of the lipid staining signal **(Figure 5C-D**, **figure 5E)**, similar to previous studies^55^. This pattern was similar in diploid and tetraploid animals and therefore could not explain the different survival phenotype of tetraploids upon cold recovery.

**Figure 5.**
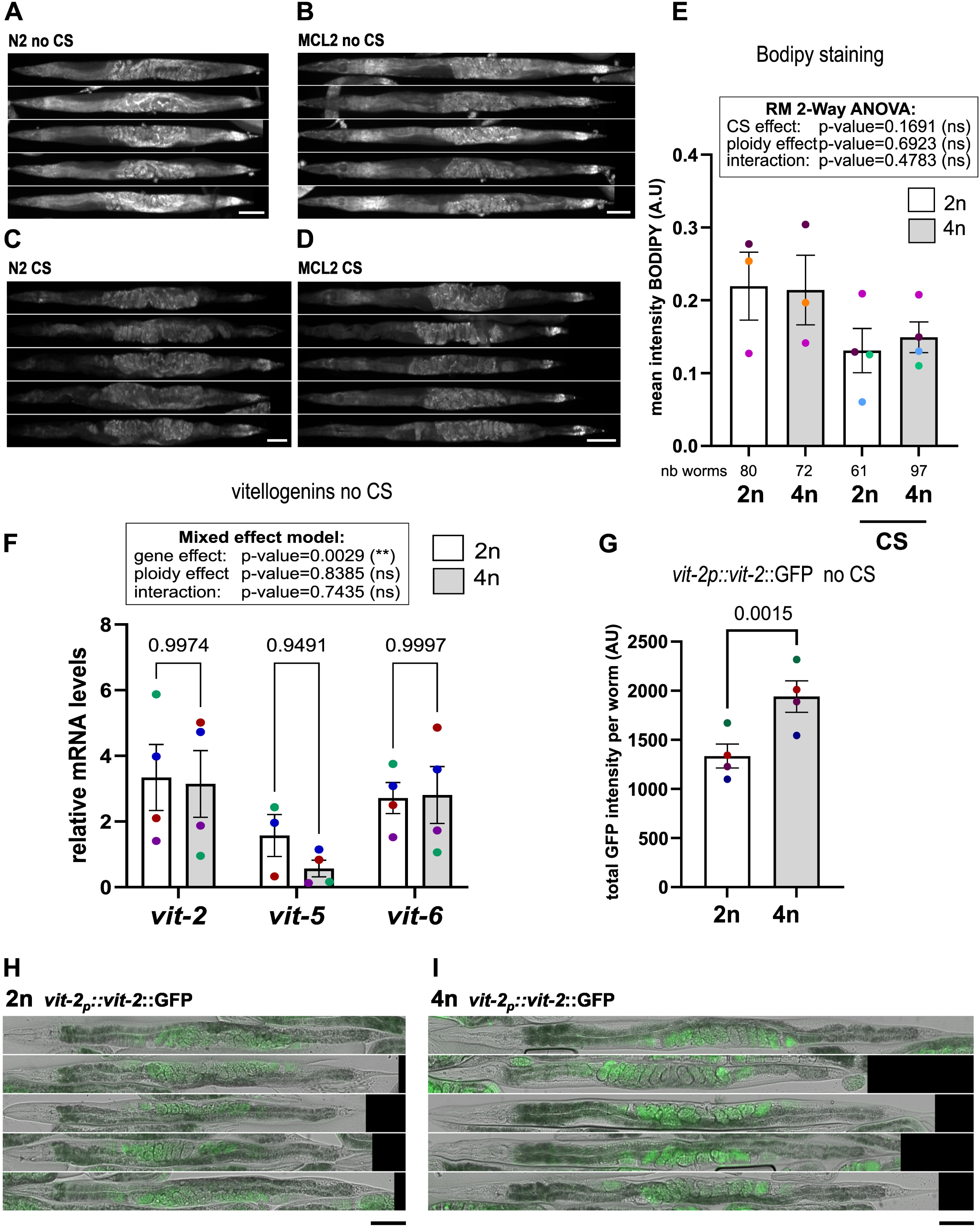
Tetraploids survival upon cold recovery is not caused by a defect in cross-tissue lipid relocalisation after cold stress, nor by a decrease in vitellogenin production and loading into embryos. **(A-D)** Diploid N2 and tetraploid MCL2 day 2 adults stained with BODIPY 493/503 in the absence of cold shock (A, B), and after 10h recovery from a severe cold shock of 4h at 0-1°C (C-D). Scalebar: 100 µm. **(E)** Mean fluorescence intensity levels per worm. RM Two-way ANOVA with uncorrected Fisher’s LSD. P-values are indicated on the graph. **(F)** relative mRNA expression levels of vitellogenin *vit-2,5,6* in day 2 adult diploids and tetraploids animals. Levels of mRNA were normalised to four housekeeping genes (*klp12, lap-2, act-1,* and *pmp-3*). Mixed model analysis with Šídák’s multiple comparison tests. The p-values for each factor are indicated on the graph. **(G-I)** Fluorescence intensity levels of the single-copy translational reporter *vit-2p::vit-2*::GFP in diploid (BCN9071) and tetraploid (MCL54) day 2 adults. Micrographs of diploid BCN9071 (H) or tetraploid (I) animals. As VIT-2::GFP was expressed exclusively in embryos, total intensity GFP levels per worm (representing levels in all embryos) were measured (G), rather than mean intensity levels (which take into account increased body size in tetraploids). Paired t-test. Colours indicate matching independent biological replicates. Error bars = SEM. Scale bar: 100 µm.

Maternal provisioning and lipid movement from the intestine to the germline in embryos are controlled by vitellogenin proteins, belonging to three families encoding yolk proteins YP170B (*vit-1-2*), YP170B (*vit-3-4-5*) and peptides YP115 and YP188 (*vit-6*)^56^. Loss of function of *vit-2* and *vit-5* are associated with increased survival upon cold recovery and inhibition of soma-to-germline lipid relocalisation following cold shock^55^. We asked whether vitellogenin levels could be decreased in tetraploids, thus increasing survival following cold shock. Nevertheless, mRNA levels of members for each of the vitellogenin families were globally unchanged in diploids and tetraploids (**Figure 5F**). We also monitored fluorescence levels from a single-copy *vit-2p::vit-2::*GFP translational reporter which revealed increased VIT-2 levels in tetraploid embryos, compared to diploids (**Figure 5G-I**). Altogether, our data suggests that the resistance of gravid adult tetraploids to severe cold shock is neither caused by a defect in cross-tissue lipid relocalisation following cold shock nor by decreased vitellogenin levels.

### Decreased induction of the *zip-10* cold-activated death program in tetraploids

Previous work has shown that, upon recovery from severe cold stress, a genetic death program is activated. Upon recovery from severe cold, the transcription factor *zip-10* is upregulated, leading to activation of several cathepsin-like proteases (including *asp-17* and *cpr-3*), and triggering organismic death (phenoptosis)^57^. We asked whether the *zip-10* cold-induced death program is inactive in tetraploids as a possible mechanism of resistance upon cold recovery.

To capture the beginning of the cold-activated transcriptional response, we monitored mRNA levels of *zip-10*, *asp-17* (*zip-10* induced target), and two other *zip-10* independent targets (*srr-6* and F53A9.5) after a short cold shock of 30 minutes. Following cold shock, all four mRNAs were induced in diploids and to a lesser extent in tetraploids for *zip-10*, *asp-17,* and *srr-6* **(Figure 6A-D).** While there was significant upregulation of all four targets upon cold recovery, the ploidy effect varied across time points, with the interaction between ploidy and time being significant for all four targets. In diploids, the maximum induction was observed at 30 minutes post cold shock, with a 15-fold induction of *zip-10* mRNA levels, about twice the previously reported levels^57^. However, in tetraploids, at 30 minutes post cold shock, *zip-10* and *asp-17* mRNA levels were respectively 2.7- and 5.8-fold less induced compared to diploids **(Figure 6A-B).** Nevertheless, there was no significant decrease at this time point for *zip-10* independent cold-activated targets *srr-6* and F53A9.1 **(Figure 6C-D).** Thus, the *zip-10* death program^57^ is also induced in our cold recovery conditions, and the activation of this program is decreased in tetraploids. We noticed, however, that the levels of the four mRNAs cited above were higher at basal level in tetraploids compared to diploids **(Figure S5B).**

**Figure 6.**
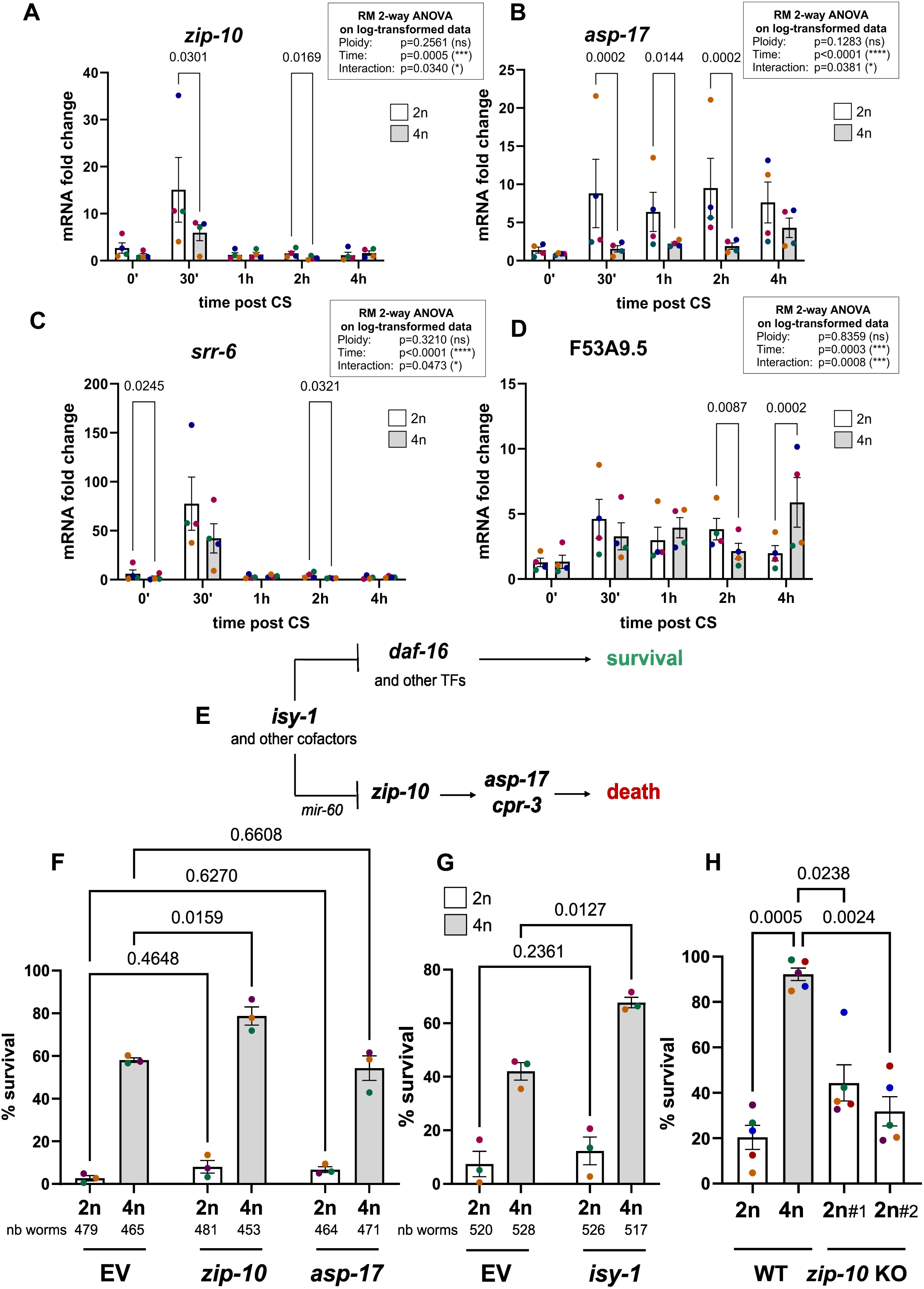
Following CS, the activation of the *zip-10* program is reduced in tetraploids and *zip-10* and *isy-1* knock-down further enhance tetraploid survival. **(A-D)** mRNA expression levels following a 30-minute cold shock (CS) of the transcription factor *zip-10* **(A)**, its target *asp-17* **(B)**, as well as cold-induced *srr-6* **(C)** and F53A5.9 **(D).** Levels of mRNA in day 2 adults were normalised to the housekeeping genes (*klp12, lap-2, act-1,* and *pmp-3*). RM Two-way ANOVA on log-transformed data (as data is log-normally distributed) with Tukey’s multiple comparison post-hoc tests were performed. For clarity, the graphs depict non-log-transformed data. For each ploidy, the mRNA levels were normalised such that levels without CS=1. The p-values for each factor of the Two-way ANOVA are indicated on the graph. For clarity, only the significant p-values of the post-hoc tests are indicated on the graph. **(E)** Schematic model of *zip-10* controlled organismic death program and the *isy-1* paradox. On one hand, the transcription factor ZIP-10 is epistatic to ISY-1 in the induction of the proteases *asp-17* and *cpr-3* upon cold recovery. However, ISY-1 also antagonizes DAF-16 and other stress transcription factors, which play a pro-survival role upon cold recovery (the *isy-1* paradox). As reported in ref^57^, after cold recovery, the primary effect is attributed to the *daf-16* branch in 2n, with *isy-1* KD providing protective effect in a *daf-16* dependent manner. Adapted from ref^57^. **(F)** Effects of *zip-10* and *asp-17* knockdown by RNA interference (RNAi) on cold recovery of day 2 adults of each ploidy. RM Two-way ANOVA with Tukey’s multiple comparison post-hoc tests were performed. P-values for main effects: ploidy effect: p=0.0020 (**), RNAi target: p=0.0028(**), interaction: p=0.0384 (*). P-values for the comparison 2n/4n for EV: p=0.0002 (***), *zip-10*: p<0.0001 (****) and *asp-17*: p=0.0003 (***). **(G)** Effect of *isy-1* knockdown by RNAi on cold recovery of day 2 adults of each ploidy. RM Two-way ANOVA with uncorrected Fisher’s LSD was performed. P-values for main effects: ploidy effect: p=0.0032 (**), RNAi target: p=0.0004 (***), interaction: p=0.0375 (*). P-values for the comparison 2n/4n for EV: p=0.0071 (**), *isy-1*: p=0.0028 (**). In (F-G), each dot represents a biological replicate including 3 technical replicates. The total number of worms assayed across all replicates is indicated on the graph below each condition. **(H)** Survival of *zip-10(ok3462)* loss of function mutants (outcrossed 4 times) upon cold recovery. RM One-way ANOVA with Geisser-Greenhouse correction: p-value=0.0001(***). Significant p-values for multiple comparisons with Tukey’s multiple comparison correction are indicated. In all graphs, error bars = SEM and colours indicated matching independent biological replicates.

We tested the effect of knocking down (KD) components of the cold-induced program **(Figure 6E)** on survival after cold recovery. RNAi against *zip-10* significantly increased the survival of tetraploids, possibly caused either by further reduction of *zip-10* levels in tetraploids or by a synergy between the *zip-10* program and the mechanisms underlying tetraploid survival upon cold recovery **(Figure 6F)**. However, *zip-10* RNAi did not significantly increase the survival of diploids in our cold stress conditions. RNAi against *zip-10* induced protease *asp-17* had no effect on tetraploid survival, despite *asp-17* RNAi knock-down efficiency being more than 80% in both diploids and tetraploids **(Figure S5C).** It is likely that knocking down additional *zip-10* targets is required to observe an effect on survival from cold recovery. We also knocked down an upstream regulator *of zip-10*: the splicing homolog factor *isy-1. Isy-1* is involved in different pathways with opposite effects on survival from cold recovery^57^ **(Figure 6E)**. i*sy-1* downregulates *zip-10,* therefore *isy-1* KD would then be expected to increase death rates. However, *isy-1* KD leads to increased survival from cold recovery in diploids (the *isy-1* paradox), via the activation of protective stress transcription factors (mainly through *daf-16*)^57^. Our data shows that *isy-1* KD significantly increase survival of tetraploids **(Figure 6G)**. This positive effect of *isy-1* on tetraploid survival is similar to the effects of *isy-1* KD in diploids reported in^57^ mainly through *daf-16*. This suggests a synergy between the *zip-10* program and mechanisms responsible for tetraploid survival upon cold recovery. It is also possible that DAF-16 and/or other stress transcription factors (TFs) are partially activated in tetraploids and that *isy-1* KD further activates those TFs.

Our data shows that *zip-10* induction upon cold recovery is reduced in tetraploids and that further reduction in *zip-10* levels by RNAi enhances survival post cold recovery. The question remains whether the survival phenotype is controlled by *zip-10* dosage, such that reduced *zip-10* levels would enable tetraploids to escape from a cold-induced death program. To determine the effects of the total absence of any *zip-10* mRNA on survival post cold recovery, we subjected diploid *zip-10(ok3462)* null mutants (outcrossed four times) to cold recovery. As expected, diploid *zip- 10(ok3462)* null mutants show a trend in increased survival post cold recovery: ∼4-5 fold higher than WT, although this is not statistically significant **(Figure 6H).** However, the survival rate of diploid null *zip-10* mutants is more than two times lower than WT tetraploids, indicating again that the strong tetraploid cold recovery survival phenotype cannot be solely explained by a decrease in *zip-10* activity. Our cold recovery survival results were different when performed on *zip-10(ok3462)* non-outcrossed mutants **(Figure S5D)**; where non-outcrossed *zip-10(ok3462)* exhibited a very high survival upon cold recovery (similar to tetraploid animals), suggesting that this strong cold recovery survival phenotype was caused by background mutations. Altogether, our data suggests that, although *zip-10* cold-induction is reduced in tetraploids, this decrease can only partially explain their survival. This indicates that additional, as yet unidentified mechanisms contribute to the ability of gravid adult tetraploids to escape cold-induced death.

## DISCUSSION

Whether autopolyploidy might be beneficial for a multicellular animal in a stressful environment is a matter of ongoing debate. In this study, we explored the consequences of induced autotetraploidy on physiology and investigated how it affects stress responses in the nematode *C. elegans*. We show, that under regular conditions, induced autotetraploidy has deleterious consequences on fitness, with decreased fertility, longevity, and developmental delay. The effects of tetraploidy on fertility recapitulate previous findings^36,58^.

In contrast to autopolyploid plants^32,37^, exposure to pathogenic bacteria did not lead to an altered response in neotetraploid *C. elegans*. Exposure to heat stress revealed a modest increase in thermorecovery, possibly linked to decreased surface/volume nuclear ratio in tetraploids, as suggested by alteration in the *hsp-16.2* DNA locus after heat shock. We did not, however, observe any consequence on the dynamics of the heat shock response. It is possible that high variability in our qRT-PCR results between independent biological replicates **(Figure 2K-M)** might mask subtle differences in heat shock kinetics patterns between animals of different ploidies. An alternative hypothesis is that the positive effect of tetraploidisation on resistance to heat stress is mediated by changes in other biological pathways, such as translation inhibition, or an increase in protein degradation.

Our study reveals for the first time that induced autotetraploidy in an animal model alters responses to temperature-related stresses, and that gravid autotetraploid *C. elegans* escape cold-induced death with potential adaptive consequences. Cytogenetic studies in the 1970-90’s reported individual occurrences of polyploidy in several nematode species, without those events being fixed in the population^59–62^. *C. elegans* are poikilotherms and cannot regulate their own body temperature. While *C. elegans* have evolved mechanisms of gradual adaptation to cold, such as adjusting membrane fluidity and metabolic cold adaptation^63,64^, sporadic tetraploidisation of some individuals within the population may diversify the outcomes from severe cold stress, with tetraploid individuals escaping cold induced death and producing more progeny of equal fitness than diploids.

To uncover the mechanisms underlying tetraploid escape from cold-induced death, we investigated known pathways involved in response to severe cold. We found that lipids are stored mainly in the intestine and embryos under regular conditions. Upon cold recovery, intestinal lipid stores are depleted, while lipids are still detected in the germline. These results differ from ref.^55^, who report an absence of lipids in embryos under regular conditions, and following cold shock an increase in the number of embryos positively stained for lipids, interpreted as intestine-to-germline lipid relocalisation. However, we repeated our experiments which produced robust results. While we do not fully understand the reasons underlying these discrepancies, it could be linked with the nature of the lipid staining dye. Altogether our study suggests that cold recovery affects lipid localisation patterns similarly between diploids and tetraploids, thus uncoupling cold-induced death from cold-induced lipid pattern changes in animals of different ploidies. In support of this, the transcriptional activity of vitellogenin lipid transporters was similar in tetraploids compared to diploids.

Our investigation of the *zip-10* cold-activated death program revealed decreased induction of the *zip-10* program in cold shocked tetraploids. However, our analysis suggests that this can only partially explain the survival phenotype of tetraploids post cold recovery. On one hand, further decreasing *zip-10* levels enhances tetraploid survival, and on the other hand, the absence of *zip-10* in diploids does not fully phenocopy the resistance phenotype of tetraploids. This indicates that additional mechanisms underly the survival of tetraploids upon cold recovery.

Metabolism is likely to play an important role in survival after cold recovery. First, raising diploid worms at 15°C (acclimatising) enables 100% survival upon cold recovery^53^. Yet, lowering growth temperature shifts metabolism to adapt to lower temperature, for instance via upregulation of fatty acid desaturase enzyme *fat-7*^64,65^. Second, we noticed lower basal survival rates upon cold recovery when the worms were fed HTT115 (40-60% survival of tetraploids, during RNAi experiments) instead of OP50 bacteria (>80% survival of tetraploids), highlighting the influence of diet and metabolism on survival from cold recovery. Alternatively, this could be linked to mild activation of innate immune response on worms fed with slightly pathogenic OP50. Several studies in other models showed an association between ploidy changes and metabolism^2,5,45,66,67^. This suggests that metabolic changes associated with WGD could underlie the survival of tetraploid adults upon cold recovery. The stress transcription factor DAF-16 could be an interesting candidate player. DAF-16 integrates signals from the environment, downstream of insulin and TOR signalling pathways, to regulate metabolism and has a protective role during cold recovery, in parallel to the *zip-10* pathway^57^. However, our data suggests synergistic effects between stress transcription factors including DAF-16 (as seen upon *isy-1* RNAi) and other mechanisms underlying the survival of tetraploid adults upon cold recovery.

Another possibility is that the escape from cold-induced death in adult tetraploids is linked to gonadal signalling and/or germ-cell proliferation. The fact that survival after cold recovery in 2n/4n differs at the gravid adult stage, but not at the L4 stage, points to a potential link with the germline. In line with this, diploid worms lacking a gonad (this study) escape cold induced death similarly to tetraploids. Furthermore, diploids, having a gonad but lacking germ cell proliferation (*glp-1* loss of function mutation) escape cold-induced death better than WT^55^. Previous work has shown that signalling from the reproductive system affects fat metabolism ^68,69^.

Other possible mechanisms could involve neuronal signalling. Indeed, cold induced death requires functional TAX-2/TAX-4 cGMP receptor signal from a subset of thermosensory neurons^55^ and the G-coupled protein receptor FSHR-1, (expressed in neurons and intestine) is also an identified mediator of cold-induced programmed death^70^. Further studies are needed to help fully uncover the mechanisms of survival to cold recovery at play in synthetic autotetraploids.

Our work provides, for the first time, evidence that autotetraploidy in animals is associated with increased stress survival, with potential adaptive implications. While this manuscript was in preparation a study was published that investigated the consequences of autotetraploidy in *C. elegans* on physiology and response to chemotherapeutic drugs^71^. Our study is, however, the first one to investigate stress responses to heat, cold and pathogens in a synthetic autotetraploid multicellular animal. Our findings point to the involvement of novel, previously unidentified mechanisms that enable neotetraploid adults to escape cold-induced death. Further research will be essential to fully elucidate the mechanisms driving tetraploid adult *C. elegans* survival following exposure to severe cold. While the specific mechanisms underlying tetraploid stress resilience, and the involvement of the gonad, and presumed germline-signalling are likely to be nematode-specific, whether the general phenomena underlying altered cell biology and metabolism may be shared across polyploids is an exciting new avenue of research.

## ACKNOWLEDGEMENTS

We are grateful to Professor Marie-Anne Félix and Professor Henrique Teotónio from Institut de Biologie de l’École Normale Supérieure (IBENS) de Paris for helpful discussions. We are thankful to Dr Carsten Kröger from the Microbiology Department at Trinity College for help with the *Pseudomonas aeruginosa* experiment. We would like to thank Dr Gavin McManus from the School of Biochemistry and Immunology Imaging facility at Trinity College, Dr Karsten Hokamp from the bioinformatics facility in the Genetics Department at Trinity College and Dr Pablo Labrador in the Genetics Department at Trinity College for all their help. We thank the *Caenorhabditis elegans* Center (CGC) for providing some strains. The CGC is funded by NIH Office of Research Infrastructure programs (P40 OD010440). This work was supported by funding from the European Research Council, grant agreement 771419, and from Research Ireland (SFI-IRC Pathway grant 22/PATH-S/10765).

## GENERAL METHODS

### *C. elegans* maintenance

Nematodes were grown on NGM plates seeded with *Escherichia coli* OP50 strain at 20**°**C unless otherwise stated, according to standard methods^72^.

### List of *C. elegans* strains

To distinguish between the genotypes of diploid and tetraploid worms, the following nomenclature was used, similar to^35^. The number of autosomal chromosomes is listed as 2A in diploid and 4A in tetraploid animals. Neotetraploid hermaphrodites can carry either 3X or 4X chromosomes^36,73^ with “allélogènes” 4A:3X siring 40% males in their progeny while “thélygènes” 4A:4X hermaphrodites exhibit 0.7% males in their progeny. All the derived tetraploid lines generated here carry 4X chromosomes, based on male frequency in their progeny.

The following strains were used: N2 (2A:2X) WT, AM140 (2A:2X) *rmIs132* [*unc-*54p::Q35::YFP], AM738 (2A:2X) *rmIs297 [pAMS66 vha-6p::Q44::YFP + rol-6(su1006) + pBluescript II],* BCN1049 (2A:2X) *crgIs1004[daf-21p::gfp; unc-119(+)]*, BCN1082 (2A:2X) *crgSi1004[pdaf-21::gfp); unc-119(+)] II at ttTi5605 x4*, BCN9071 (2A:2X) *crg9070[vit-2p::vit-2::gfp]) X*, CB1331 (2A:2X) *unc-54(e1301)* I temperature sensitive, EJ1171 (2A:2X) *gem-1(bc364); gon-2(q388)* temperature sensitive, GW615 (2A:2X) *gwSi3* [*hsp-16.2::wmCherry; 256x LacO; unc-119(+)]; gwIs39[baf-1::gfp-LacI::let-858 3’UTR; vit-5::gfp*] III; *unc-119(ed3)* III; RB2499 (2A:2X) *zip-10(ok3462)* non outcrossed. The RB2499 strain was genotyped using the following primers: *zip-10 (ok3462)* For 5’-GCACAACTCGGGTGCTCATA and *zip-10 (ok3462)* Rev 5’-AAGAAACGAGGTGGGGATGG. The *ok3462* allele is a 661 nucleotides (nt) deletion starting 71 nt before the start codon and ending 40 nt after the stop codon and with the insertion of the following 10 nt indel: AATTAAAAAA. With the entire 549 nt coding sequence removed, *ok3462* is a loss of function allele of *zip-10*.

The following strains were generated. Tetraploid derivative of N2: MCL1 (4A:4X) WT line #1, MCL2 (4A:4X) WT line #2. Tetraploid derivative of AM140: MCL6 (4A:4X) *rmIs132* [*unc-54p*::Q35::YFP]. Tetraploid derivative of GW615: MCL7 (4A:4X) *gwSi3* [*hsp-16.2::wmCherry; 256x LacO; unc-119(+)]; gwIs39[baf-1::gfp-LacI::let-858 3’UTR; vit-5::gfp*] III; *unc-119(ed3)* III. Tetraploid derivatives of AM738: MCL20 (4A:4X) *rmIs297* [pAMS66 *vha-6p*::Q44::YFP + *rol-6(su1006)* + pBluescript II] line #1, and MCL21 (4A:4X) *rmIs297* [pAMS66 *vha-6p*::Q44::YFP + *rol-6(su1006)* + pBluescript II] line #2. Tetraploid derivative of EJ1171: MCL22 (4A:4X) *gem-1(bc364); gon-2(q388)* temperature sensitive. Tetraploid derivatives of CB1331: MCL25 (4A:4X) *unc-54(e1301)* I temperature sensitive line #1, and MCL26 (4A:4X) *unc-54(e1301)* I temperature sensitive line #2. Tetraploid derivatives of BCN1049: MCL11 (4A:2X) *crgIs1004[daf-21p::gfp; unc-119(+)].* Tetraploid derivatives of BCN1082: MCL36 (4A:2X) *crgSi1004[daf-21p::gfp); unc-119(+)] II line#2* and MCL38 (4A:2X) *crgSi1004[daf-21p::gfp); unc-119(+)] II line#1.* Tetraploid derivatives of BCN9071: MCL54: (4A:2X) *crg9070[vit-2p::vit-2::gfp]) X.* Diploid outcross (four times) of RB2499: MCL66 *zip-10(ok3462) x4 line#1*, and MCL67 *zip-10(ok3462) x4* line#2.

### Generation of tetraploid *C. elegans* strains

Autotetraploid strains were generated by RNA interference of the cohesin component complex *rec-8* for two generations and selection of longer worms in the F2 progeny^35^. Longer worms were passed for a few generations, typically 3-10, before stable lines producing only tetraploids were obtained (Occasionally, diploid revertant worms were observed). The ploidy status of derived tetraploid strains was confirmed by visually monitoring the number of chromosomes in −1 oocytes in DAPI stained animals using a Zeiss SP8 confocal microscope similarly to ref^35^.

### Worm synchronization and staging

Worms were synchronized by egg-laying of several 6 cm plates each containing 20-30 gravid adult worms for diploids and about 60 gravid adults for tetraploids. Gravid adults were then removed, and the progeny were assayed at the indicated time post synchronization. For experiments involving the gonadless mutant *gon-2(q388);gem-1(bc364)*, gravid adults were synchronized for 12-14h at the permissive temperature (15°C). Gravid adults were removed, and nematodes were then switched to the restrictive temperature (25°C). Nematodes were then selected at late L4 larval stage. For experiments at specific stages (L4 or Day 2 adults), worms were selected based on the late L4 stage^74^ and were assayed 24h later (as day 2 of adulthood).

### Fertility assay

About 20 worms per condition were singled out in 12-well plates seeded with 50-μL OP50 at the L4 stage. Each day, all animals were passed onto new 12-well plates. The F1 progeny laid by each individual P0 worm was scored 2 to 3 days after the parent worm had been transferred to the well, when the F1s were either L4 or adults. For fertility experiment after cold shock, Gravid day 2 adult cold-shocked worms were singled out immediately after the end of cold shock in 12-well plates. Their progeny was assayed 3 days later and the P0 parent worm was transferred to a second well if it was alive 72h post cold shock, and its progeny was scored 3 days later.

### Embryonic lethality assay and developmental delay

Gravid animals were allowed to lay eggs for two hours. A minimum of 100 embryos per condition were transferred onto a new plate, just outside of the bacterial lawn. The next day, embryos that did not hatch were scored as dead. At 65h post egg-laying synchronization at 20°C, the stage of each individual animal was determined. A developmental index was calculated by attributing a score from 1 to 6 to the successive stages L1, L2, L3, L4, young adult stage and gravid adult.

### Lifespan

Lifespan assays were performed at either 20°C or 25°C as previously described^75^. Worms were synchronized by egg laying within 2 hours, as described above. A total of 50 hermaphrodites were cultured on each 6-cm NGM petri dish, seeded with 250 µLOP50. Animals were transferred to a fresh plate every 1 to 2 days until the cessation of progeny production and every 2 to 3 thereafter. Animals were scored every 2 to 3 days, and recorded as dead if they showed no spontaneous movement or response when probed on the nose. Animals dead from internal hatching (“bagging”), extruded intestine, and from desiccation on the side of the plate were censored.

### Resistance to pathogenic bacteria

Experiments were performed at 25°C, according to^40,41^. Fresh cultures of *Pseudomonas aeruginosa* PAO1 (<16h culture) were used to seed NGM plates. Plates were seeded with 50 µL bacterial culture by spreading on the totality of the NGM surface. Plates were left to dry for 48h before transferring the worms. Only fresh plates (no more than two days at room temperature) were used. For experiments involving N2 and MCL2, 40 µL of FUdR (5-Fluoro-2’-deoxyuridine) at 100mg/mL was pipetted onto each assay plate about 30 min-1h hour before transferring the worms on it, to prevent internal hatching, similar as in (Kirienko et al. 2014). Worms were fed OP50 bacteria until late L4 stage, when they were transferred on plates containing pathogenic bacteria. Not more than 40 worms were transferred onto each plate and worms were assayed every day.

### Heat-shock

NGM plates were double sealed with parafilm, and heat shocked in a water bath. Unless otherwise stated, heat shock was 30 minutes at 34°C. Worms were harvested immediately after heat shock unless stated otherwise.

### Thermorecovery assay

The thermotolerance assay was performed as described previously^76^, except that heat-shock was 4h at 36**°**C. About 60-100 synchronized animals (late L4 stage) were picked onto a 6 cm NGM plate. Animals were then allowed to recover overnight at 20**°**C or 25**°**C. Animals were scored the next day at about 20h after the end of heat-shock. Animals were transferred onto a new plate and were counted as alive when they were either moving on the plate, or at least able to move their nose when poked with a pick.

### Motility assay in *unc-54(e1301)* temperature sensitive mutants

Worms were raised at 15°C and transferred to 25°C at the L4 stage for experiments at the restrictive temperature. The motility was assayed 15h later. A circle of 1 cm diameter was drawn around the center of the lid of a 6 cm plate, which was placed under an NGM plate were OP50 bacteria were seeded at the periphery. Worms were transferred at the center of the 1 cm diameter circle and their position was scored after 2 minutes, and classified in 3 zones: outside the circle, inside the circle or close to the center of the circle.

### Cold recovery assay

At least 60 to 100 worms were transferred onto a fresh seeded NGM plate per condition. Plates were sealed with parafilm and buried upside down in a Styrofoam box containing ice. The ice box was placed in a 4°C cold room for 4 hours. After the end of cold shock, plates were placed back in the incubator at the desired temperature for 20h, before scoring of survival. The temperature during cold shock **(Figure S5A)** was monitored using a thermocouple thermometer HH-613 Thermosense with a probe placed at the surface of the NGM in a 6 cm Petri dish placed inside a Styrofoam filled with ice.

### Quantitative real-time PCR

To monitor steady-state mRNA levels, we handpicked a pool of about 50 animals per condition in 20 μL of RNase free water. 500 µL of TRIzol was added and samples were processed as described in^77^. Reverse transcription was carried out using the Revert Aid First Strand synthesis kit from Thermo Scientific according to the manufacturer’s instructions using oligo dT primers. The concentration of cDNA was monitored on a Nanodrop. Measurements of mRNA levels were obtained by qRT-PCR on a Lightcycler 480 (Roche). The amount of cDNA was quantified using the delta Ct quantification method, assuming 100% PCR efficiency for every couple of primers. For each couple of primers, the PCR efficiency was calculated by running a standard curve on a dilution series. Validated couples of primers had a PCR efficiency between 90 and 113% with R2 *>* 0.98. Expression levels of steady-state mRNA were calculated using the ΔΔCt method.

### Determination of housekeeping genes for qRT-PCR normalization

Primer sets (listed in Table S1) were designed to span exon-exon junctions (using NCBI Primer Blast software), and subsequently blasted against the *C. elegans* genome to test for off-target complementarity. It is crucial to verify the stability of the candidate housekeeping genes for normalization when assessing gene expression between different ploidies, to ensure that the expression of those genes relative to the others is not affected by the change of ploidy. Each target mRNA was normalized to the average of the optimal number of the most stable housekeeping genes determined using the geNorm algorithm^78^, which establishes a hierarchy of stable genes (from a set of minimum 8 candidates) from pairwise comparisons of expression levels across all conditions (minimum 10 different samples).We analysed a set of 12 candidates genes (Y45FIOD.5, *pmp-3*, *act-1*, *tsn1*, *klp-12*, *unc-16*, *lap-2*, *ife-1*, *ire-1*, *cdc-42*, *gpd-2* and *ama-1*) on 10 sets of 4 identical conditions (N2 and MCL2 strains, at both L4 and D2 stage). Three separate analyses were conducted, using N2 and MCL2 strain at L4 stage, D2 stage or both L4 and D2 stage. For gene expression analysis performed at the L4 stage only, the GeNorm algorithm recommended to normalize to the average of the five most stable housekeeping genes: respectively Y45F10D.4, *pmp-3*, *lap-2, klp-12,* and *act-1*. However, we show that there was no difference in normalizing expression data at L4 stage to the average of the top three most stable housekeeping genes only (Y45F10D.4, *pmp-3* and *lap-2*), as seen in **Figure 2 K-N**, compared to normalizing to the average of the five most stable housekeeping genes (**Figure S1**). Therefore, we kept using the three most stable housekeeping genes for normalizing gene expression data at the L4 stage. For gene expression analysis between L4 and day 2 adults, a second geNorm analysis was performed on genes expressed in the soma only^79^, to avoid potential artefacts coming from differences in the germline between diploids and tetraploids. Expression levels were normalized to the average of Y45F10D.4, *pmp-3* and *lap-2* for qRT-PCR analysis at L4 and Day 2 adult stage. For gene expression analysis performed at the D2 stage only, expression levels of target genes were normalised to the average of *pmp-3*, *lap-2*, *act-1* and *klp-12*.

### Fluorescence microscopy and aggregates quantification

Animals were paralyzed in 3mM Levamisole diluted in M9 and mounted on a 2% agarose pad. Fluorescent images were taken on an epifluorescence Olympus IX81 microscope. At objective 10 X, unless otherwise stated. To quantify aggregates in worms carrying Q35:;YFP or Q44::YFP, the number of aggregates was counted for each individual worm on the original images and normalized to the length of each worm (measured using FIJI/Image J), to account for body size increases in tetraploid lines. For visualization purpose, worms were straightened in some cases, using the FIJI/ImageJ macro Worm-align^80^. Images of worms carrying the single copy transgene *hsp-16*p-lacO/UASp::LacI were taken on a Zeiss SP8 confocal microscope at objective 63X. Z-stacks were acquired using a step of 0.2 µm.

### Monitoring of *hsp-16* DNA locus within the nucleus

The *hsp-16.2* DNA locus was visualized using the in vivo tracking system developed by^44^ using animals carrying the single copy transgene *gwSi3* [*hsp-16.2::wmCherry; 256x LacO; unc-119(+)]* together with the multicopy transgene *gwIs39[baf-1::gfp-LacI::let-858 3’UTR; vit-5::gfp]*, either in diploid (GW615) or in a tetraploid (MCL7) context. Gravid adults of each genetic background were dissected, and embryos were mounted on a slide and immediately imaged on a confocal microscope. Z-stacks of early live embryos (not more than ∼50 cell stage) were acquired on a confocal microscope. Heat-shock was performed next to the confocal microscope by warming the glass slide containing live embryos on a heating block for 10 minutes at 34°C. Heat shocked embryos were immediately imaged at the end of heat shock. Measuring *hsp-16.2* DNA loci position in the nucleus was performed according to^49^. Z-stacks containing images were visualized using the image analysis software Imaris (BITPLANE, Oxford Instruments). The *hsp-16.2* loci dots were selected and coloured differently from the diffuse nuclear GFP, based on GFP intensity thresholds (**Figure 2C**). The shortest distance to the nuclear envelope was measured for each *hsp-16.2* locus dot, together with the diameter of the nuclear plane in 2D images. The position of the foci was then classified in 3 zones of equal surface area, with zone 1 being the closest to the nuclear envelope (**Figure 2F**).

### Fat staining

A minimum of 100 worms per condition were selected either at the late L4 stage or assayed 24h later (day 2 adults). Worms were fixated with 60% isopropanol. Lipid staining with BODIPY 493/503 was performed according to ref.^80^. Animals were mounted and imaged the same day or the next day on an upright Olympus IX81 epifluorescence microscope, using identical intensity settings per experiment. Lipid staining with Oil Red O (ORO) was carried out similarly to ref^81^. Worms were mounted and imaged the same day on an Olympus BX81 microscope equipped with an Olympus DP73 colour camera at objective 10X. The same exposure was used across all conditions within an experiment and images were saved as a tiff file. Raw images were processed using the ImageJ/FIJI, to subtract the background, convert to greyscale and threshold the outline of worm bodies, as described in ref^82^. The levels of BODIPY or ORO were quantified in between 10 and 40 individuals per condition per independent biological replicate. Quantification methods included Worm-align and worm_CP, which utilised ImageJ/FIJI and Cell Profiler for analysis^80^.

### RNA interference

RNA interference (RNAi) experiments were performed as described in^83^ on fresh NGM plates with 1mM IPTG and 100 µg/mL Ampicillin, seeded with bacterial cultures grown overnight with 50 µg/mL Ampicillin. All RNAi clones were sequenced prior to the experiments. Diploid and tetraploid gravid adults were grown on RNAi plates and their progeny was transferred onto fresh RNAi plates at the late L4 stage. Animals were assayed 24h later at day 2 of adulthood. However, as RNAi against *isy-1* during development was lethal, we grew parental gravid adults 2n/4n on control (L4440 empty vector). Their progeny was picked at the late L4 stage and transferred onto fresh *isy-1* RNAi plates. Worms were assayed for cold recovery 24h later at the day 2 adult stage. During cold recovery assays on EV or candidate RNAi targets, we observed more variability in survival rates, even in the control conditions, suggesting that bacterial feeding with HT115 bacteria, compared with OP50 affects recovery from severe cold shock. Therefore, we performed RNAi with three technical replicates for each condition tested in each individual biological replicate. The efficiency of RNAi knock-down was determined for *asp-17* RNAi by measuring *asp-17* mRNA levels by qRT-PCR on EV or *asp-17* RNAi. We were unable to determine *isy-1* or *zip-10* knock down efficiency in a similar manner as it the dsRNA targeting *zip-10* or *isy-1* covered the entirety of the coding region and we could not design qRT-PCR primers specific for the endogenous mRNA.

### Statistical analysis

Statistical analysis was performed using GraphPad PRISM version 10. In most experiments, statistics were performed on the means from at least three independent biological replicates, each comprising at least 30 individuals. Conditions of normality and variance homogeneity was verified using QQ plots and homoscedasticity plots.

## Notes

### Competing Interest Statement

The authors have declared no competing interest.

### Summary of Updates

revised to add the ORCID number of one of the authors (Clement verdier)

## REFERENCES

1. Van de Peer, Y., Mizrachi, E. & Marchal, K. The evolutionary significance of polyploidy. Nat. Rev. Genet. 18, 411–424 (2017).

2. Schvarzstein, M., Alam, F., Toure, M. & Yanowitz, J. L. An emerging animal model for querying the role of whole genome duplication in development, evolution, and disease Polyploidization. 1–24 (2023).

3. Li, T.-N. et al. Intrahepatic hepatitis B virus large surface antigen induces hepatocyte hyperploidy via failure of cytokinesis. J. Pathol. 245, 502–513 (2018).

4. Losick, V. P., Fox, D. T. & Spradling, A. C. Polyploidization and cell fusion contribute to wound healing in the adult Drosophila epithelium. Curr. Biol. 23, 2224–2232 (2013).

5. Sladky, V. C., Eichin, F., Reiberger, T. & Villunger, A. Polyploidy control in hepatic health and disease. J. Hepatol. 75, 1177–1191 (2021).

6. Lin, Y.-H. et al. Mice with increased numbers of polyploid hepatocytes maintain regenerative capacity but develop fewer hepatocellular carcinomas following chronic liver injury. Gastroenterology 158, 1698–1712.e14 (2020).

7. Was, H. et al. Polyploidy formation in cancer cells: How a Trojan horse is born. Semin. Cancer Biol. 81, 24–36 (2022).

8. Wolfe, K. H. Origin of the yeast whole-genome duplication. PLoS Biol. 13, 1–7 (2015).

9. Van de Peer, Y., Ashman, T. L., Soltis, P. S. & Soltis, D. E. Polyploidy: an evolutionary and ecological force in stressful times. Plant Cell 33, 11–26 (2021).

10. Chen, H., Almeida-Silva, F., Logghe, G., Bonte, D. & Van de Peer, Y. The rise of polyploids during environmental catastrophes. bioRxiv (2024) doi:10.1101/2024.11.22.624806.

11. Gregory, T. R. & Mable, B. K. Polyploidy in Animals. in The Evolution of the Genome 427– 517 (Elsevier, 2005).

12. Otto, S. P. & Whitton, J. Polyploid incidence and evolution. Annu. Rev. Genet. 34, 401– 437 (2000).

13. McLysaght, A., Hokamp, K. & Wolfe, K. H. Extensive genomic duplication during early chordate evolution. Nat. Genet. 31, 200–204 (2002).

14. Dehal, P. & Boore, J. L. Two rounds of whole genome duplication in the ancestral vertebrate. PLoS Biol. 3, e314 (2005).

15. Nakatani, Y. et al. Reconstruction of proto-vertebrate, proto-cyclostome and proto-gnathostome genomes provides new insights into early vertebrate evolution. Nat. Commun. 12, 4489 (2021).

16. Simakov, O. et al. Deeply conserved synteny resolves early events in vertebrate evolution. *Nat*. Ecol. Evol. 4, 820–830 (2020).

17. Jaillon, O. et al. Genome duplication in the teleost fish Tetraodon nigroviridis reveals the early vertebrate proto-karyotype. Nature 431, 946–957 (2004).

18. Barker, M. S., Arrigo, N., Baniaga, A. E., Li, Z. & Levin, D. A. On the relative abundance of autopolyploids and allopolyploids. New Phytol. 210, 391–398 (2016).

19. Redmond, A. K., Casey, D., Gundappa, M. K., Macqueen, D. J. & McLysaght, A. Independent rediploidization masks shared whole genome duplication in the sturgeon-paddlefish ancestor. Nat. Commun. 14, 2879 (2023).

20. Lien, S. et al. The Atlantic salmon genome provides insights into rediploidization. Nature 533, 200–205 (2016).

21. Baduel, P., Bray, S., Vallejo-Marin, M., Kolář, F. & Yant, L. The “Polyploid Hop”: Shifting challenges and opportunities over the evolutionary lifespan of genome duplications. Front. Ecol. Evol. 6, 1–19 (2018).

22. Bomblies, K. Learning to tango with four (or more): the molecular basis of adaptation to polyploid meiosis. Plant Reprod. 36, 107–124 (2023).

23. Westermann, J., Srikant, T., Gonzalo, A., Tan, H. S. & Bomblies, K. Defective pollen tube tip growth induces neo-polyploid infertility. Science 383, eadh0755 (2024).

24. Yant, L. et al. Meiotic adaptation to genome duplication in Arabidopsis arenosa. Curr. Biol. 23, 2151–2156 (2013).

25. Bray, S. M. et al. Kinetochore and ionomic adaptation to whole genome duplication. (2023) doi:10.1101/2023.09.27.559727.

26. Selmecki, A. M. et al. Polyploidy can drive rapid adaptation in yeast. Nature 519, 349– 352 (2015).

27. Lu, Y.-J., Swamy, K. B. S. & Leu, J.-Y. Experimental evolution reveals interplay between Sch9 and polyploid stability in yeast. PLoS Genet. 12, e1006409 (2016).

28. Ebadi, M. et al. The duplication of genomes and genetic networks and its potential for evolutionary adaptation and survival during environmental turmoil. Proc. Natl. Acad. Sci. U. S. A. 120, e2307289120 (2023).

29. Mable, B. K., Alexandrou, M. A. & Taylor, M. I. Genome duplication in amphibians and fish: An extended synthesis. J. Zool. (1987) 284, 151–182 (2011).

30. Novikova, P. Y. et al. Polyploidy breaks speciation barriers in Australian burrowing frogs Neobatrachus. PLoS Genet. 16, e1008769 (2020).

31. del Pozo, J. C. & Ramirez-Parra, E. Deciphering the molecular bases for drought tolerance in Arabidopsis autotetraploids. Plant Cell Environ. 37, 2722–2737 (2014).

32. Mehlferber, E. C. et al. Polyploidy and microbiome associations mediate similar responses to pathogens in Arabidopsis. Curr. Biol. 32, 2719–2729.e5 (2022).

33. Wang, W., He, Y., Cao, Z. & Deng, Z. Induction of tetraploids in impatiens (Impatiens walleriana) and characterization of their changes in morphology and resistance to downy mildew. HortScience 53, 925–931 (2018).

34. Harari, Y., Ram, Y., Rappoport, N., Hadany, L. & Kupiec, M. Spontaneous changes in ploidy are common in yeast. Curr. Biol. 28, 825–835.e4 (2018).

35. Clarke, E. K., Rivera Gomez, K. A., Mustachi, Z., Murph, M. C. & Schvarzstein, M. Manipulation of ploidy in Caenorhabditis elegans. J. Vis. Exp. (2018) doi:10.3791/57296.

36. Madl, J. E. & Herman, R. K. Polyploids and sex determination in Caenorhabditis elegans. Genetics 93, 393–402 (1979).

37. Saei, A. et al. The status of Pseudomonas syringae pv. actinidiae (Psa) in the New Zealand kiwifruit breeding programme in relation to ploidy level. in Acta Horticulturae vol. 1218 (2018).

38. Oswald, B. P. & Nuismer, S. L. Neopolyploidy and pathogen resistance. Philos. Trans. R. Soc. Lond. B Biol. Sci. 274, 2393–2397 (2007).

39. Zhou, L. & Gui, J. Natural and artificial polyploids in aquaculture. Aquaculture and Fisheries vol. 2 Preprint at 10.1016/j.aaf.2017.04.003 (2017).

40. Kirienko, N. V., Cezairliyan, B. O., Ausubel, F. M. & Powell, J. R. Pseudomonas aeruginosa PA14 pathogenesis in Caenorhabditis elegans. Methods Mol. Biol. 1149, 653–669 (2014).

41. Scott, E., Holden-Dye, L., O’Connor, V. & Wand, M. E. Intra strain variation of the effects of Gram-negative ESKAPE pathogens on intestinal colonization, host viability, and host response in the model organism Caenorhabditis elegans. Front. Microbiol. 10, 3113 (2019).

42. Kemp, B. J., Church, D. L., Hatzold, J., Conradt, B. & Lambie, E. J. Gem-1 encodes an SLC16 monocarboxylate transporter-related protein that functions in parallel to the gon-2 TRPM channel during gonad development in Caenorhabditis elegans. Genetics 181, 581–591 (2009).

43. Lindquist, S. THE HEAT-SHOCK RESPONSE. www.annualreviews.org (1986).

44. Rohner, S. et al. Promoter- and RNA polymerase II-dependent hsp-16 gene association with nuclear pores in Caenorhabditis elegans. J. Cell Biol. 200, 589–604 (2013).

45. Doyle, J. J. & Coate, J. E. Polyploidy, the nucleotype, and novelty: The impact of genome doubling on the biology of the cell. Int. J. Plant Sci. 180, 1–52 (2019).

46. Frawley, L. E. & Orr-Weaver, T. L. Polyploidy. Curr. Biol. 25, R353–8 (2015).

47. Buntrock, L., Marec, F., Krueger, S. & Traut, W. Organ growth without cell division: somatic polyploidy in a moth, Ephestia kuehniella. Genome 55, 755–763 (2012).

48. Bourdon, M. et al. Evidence for karyoplasmic homeostasis during endoreduplication and a ploidy-dependent increase in gene transcription during tomato fruit growth. Development 139, 3817–3826 (2012).

49. Meister, P., Gehlen, L. R., Varela, E., Kalck, V. & Gasser, S. M. Visualizing Yeast Chromosomes and Nuclear Architecture. vol. 470 535–567 (Elsevier Inc, 2010).

50. Syngelaki, E., Paetzold, C. & Hörandl, E. Gene expression profiles suggest a better cold acclimation of polyploids in the alpine species ranunculus kuepferi (Ranunculaceae). Genes (Basel) 12, (2021).

51. Jiang, Y. et al. Polyploidization of Plumbago auriculata Lam. in vitro and its characterization including cold tolerance. Plant Cell Tissue Organ Cult. 140, 315–325 (2020).

52. Deng, B. et al. Antioxidant response to drought, cold and nutrient stress in two ploidy levels of tobacco plants: low resource requirement confers polytolerance in polyploids? Plant Growth Regul. 66, 37–47 (2012).

53. Ohta, A., Ujisawa, T., Sonoda, S. & Kuhara, A. Light and pheromone-sensing neurons regulates cold habituation through insulin signalling in Caenorhabditis elegans. Nat. Commun. 5, (2014).

54. Okahata, M., Motomura, H., Ohta, A. & Kuhara, A. Molecular physiology regulating cold tolerance and acclimation of Caenorhabditis elegans. Proc. Jpn. Acad. Ser. B Phys. Biol. Sci. 98, 126–139 (2022).

55. Gulyas, L. & Powell, J. R. Cold shock induces a terminal investment reproductive response in C. elegans. Sci. Rep. 12, 1338 (2022).

56. Perez, M. F. & Lehner, B. Vitellogenins -Yolk Gene Function and Regulation in Caenorhabditis elegans. Front. Physiol. 10, 1067 (2019).

57. Jiang, W. et al. A genetic program mediates cold-warming response and promotes stress-induced phenoptosis in C. elegans. Elife 7, (2018).

58. Misare, K. R., et al. The consequences of tetraploidy on Caenorhabditis elegans physiology and sensitivity to chemotherapeutics. bioRxivorg (2023) doi:10.1101/2023.06.06.543785.

59. Triantaphyllou, A. C. & Hirschmann, H. Morphological comparison of members of the Heterodera trifolii species complex 1). Nematologica 25, 458–481 (1979).

60. Triantaphyllou, A. C. Further studies on the role of polyploidy in the evolution of meloidogyne. J. Nematol. 23, 249–253 (1991).

61. Triantaphyllou, A. C. & Riggs, R. D. Polyploidy in an Amphimictic Population of Heterodera glycines. J. Nematol. 11, 371–376 (1979).

62. Dalmasso, A. & Younes, T. Oogenese et embryogenese chez Xiphimena index (Nematoda: Dorylaimida). Annales de Zoolologie Ecologie Animale 1, 265–279 (1969).

63. Wu, G., Baumeister, R. & Heimbucher, T. Molecular mechanisms of lipid-based metabolic adaptation strategies in response to cold. Cells 12, (2023).

64. Murray, P., Hayward, S. A. L., Govan, G. G., Gracey, A. Y. & Cossins, A. R. An Explicit Test of the Phospholipid Saturation Hypothesis of Acquired Cold Tolerance in Caenorhabditis Elegans. www.pnas.org/cgi/content/full/ (2007).

65. Gómez-Orte, E. et al. Effect of the diet type and temperature on the*C. elegans*transcriptome. Oncotarget 9, 9556–9571 (2018).

66. Cadart, C., Bartz, J., Oaks, G., Liu, M. Z. & Heald, R. Polyploidy in Xenopus lowers metabolic rate by decreasing total cell surface area. Curr. Biol. 33, 1744–1752.e7 (2023).

67. Schoenfelder, K. P. & Fox, D. T. The expanding implications of polyploidy. Journal of Cell Biology 209, 485–491 (2015).

68. Shemesh, N., Shai, N. & Ben-Zvi, A. Germline stem cell arrest inhibits the collapse of somatic proteostasis early in Caenorhabditis elegans adulthood. Aging Cell 12, 814–822 (2013).

69. Antebi, A. Regulation of longevity by the reproductive system. Exp. Gerontol. 48, 596– 602 (2013).

70. Wang, C., Long, Y., Wang, B., Zhang, C. & Ma, D. K. GPCR signaling regulates severe stress-induced organismic death in Caenorhabditis elegans. Aging Cell 22, 1–12 (2023).

71. Misare, K. R. et al. The consequences of tetraploidy on Caenorhabditis elegans physiology and sensitivity to chemotherapeutics. Sci. Rep. 13, 18125 (2023).

72. Brenner, S. The Genetics of Caenorhabditis elegans. Genetics 77, 71–94 (1974).

73. Nigon, V. Effets de la polyplo\“idie chez un nématode libre. C. R. Hebd. Seances Acad. Sci. 228, (1949).

74. Mok, D. Z. L., Sternberg, P. W. & Inoue, T. Morphologically defined sub-stages of C. Elegans vulval development in the fourth larval stage. BMC Dev. Biol. 15, (2015).

75. Morley, J. F. & Morimoto, R. I. Regulation of longevity in*Caenorhabditis elegans*by heat shock factor and molecular chaperones. Mol. Biol. Cell 15, 657–664 (2004).

76. Labbadia, J. & Morimoto, R. I. Repression of the Heat Shock Response Is a Programmed Event at the Onset of Reproduction. Mol. Cell 59, (2015).

77. He, F. Total RNA Extraction from C. elegans. Bio Protoc. 1, (2011).

78. Vandesompele, J., et al. Accurate normalization of real-time quantitative RT-PCR data by geometric averaging of multiple internal control genes. (2002).

79. Knutson, A. K., Egelhofer, T., Rechtsteiner, A. & Strome, S. Germ granules prevent accumulation of somatic transcripts in the adult Caenorhabditis elegans germline. Genetics 206, 163–178 (2017).

80. Okkenhaug, H., Chauve, L., Masoudzadeh, F., Okkenhaug, L. & Casanueva, O. Worm-align and worm_CP, two open-source pipelines for straightening and quantification of fluorescence image data obtained from Caenorhabditis elegans. J. Vis. Exp. 2020, 1–13 (2020).

81. Stuhr, N. L. et al. Rapid lipid quantification in Caenorhabditis elegans by oil Red O and Nile Red staining. Bio Protoc. 12, e4340 (2022).

82. Han, S. et al. Mono-unsaturated fatty acids link H3K4me3 modifiers to C. elegans lifespan. Nature 544, 185–190 (2017).

83. Kamath, R. S. & Ahringer, J. Genome-wide RNAi screening in Caenorhabditis elegans. Methods 30, 313–321 (2003).

