## Supplementary material for "Evidence for increased stress resistance due to polyploidy from synthetic autotetraploid *Caenorhabditis elegans*": Suplemental information

### LIST OF SUPPLEMENTARY INFORMATION

**Figure S1-**Similar mRNA expression levels of highly inducible molecular chaperones normalized to either 3 or 5 housekeeping genes in L4 stage diploid or tetraploid animals.

**Figure S2-**Contrasting results obtained using different proteostasis sensors suggest a change in the behaviour of multicopy transgenes in neotetraploid animals.

**Figure S3-**Tetraploidy differentially affects multicopy and single-copy transgenes.

**Figure S4-** ORO staining shows similar levels of neutral lipids in diploids and tetraploids under regular conditions.

**Figure S5-**Data associated with Figure 3 and figure 6. Temperature during CS. Survival of cold shocked P0 at 72h post cold shock. Expression levels of cold-recovery induced mRNAs are upregulated under basal conditions in tetraploids. RNAi knock-down efficiency for *asp-17.* Survival upon CR in *zip-10(ok3462)* non-outcrossed mutant.

**Table S1**- Primers used in this study.

**Table S2**- Survival of diploids and neotetraploids exposed to *P. aeruginosa*.

**Table S3**- Lifespan of diploids and neotetraploids raised at 20°C or 25°C.

### SUPPLEMENTARY FIGURES

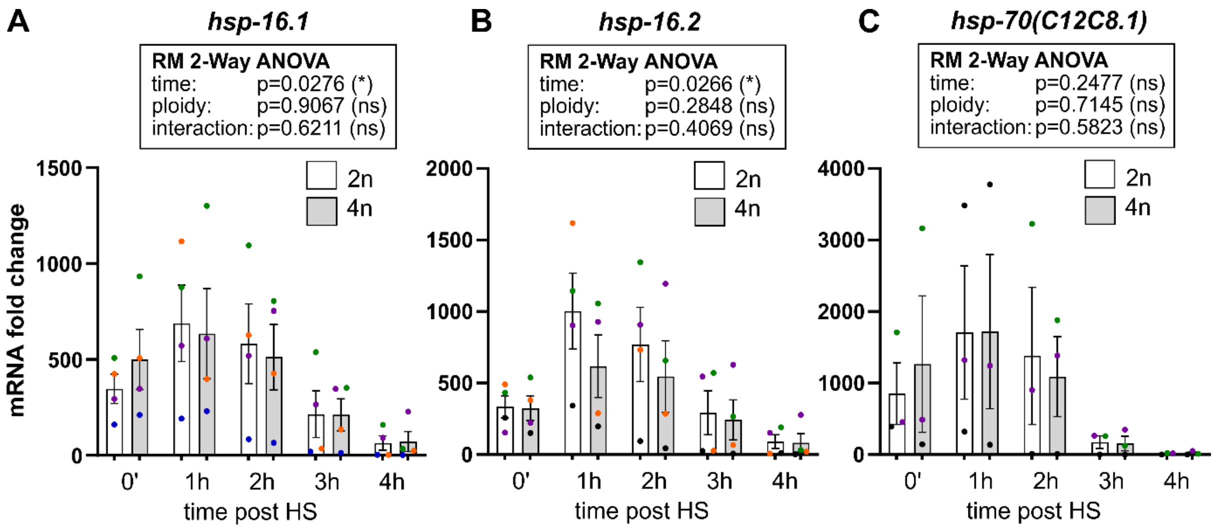


**Figure S1- Similar mRNA expression levels of highly inducible molecular chaperones normalized to either 3 or 5 housekeeping genes in L4 stage diploid or tetraploid animals.** L4 larvae were exposed to a 30-minute heat shock at 34°C. Relative mRNA expression levels of *hsp-16.1* (A), *hsp-16.2* (B) and *hsp-70(C12C8.1)* (C). As detailed in materials and methods the GeNorm algorithm[72] recommends normalizing to the five most stable housekeeping genes when measuring mRNA levels at the L4 stage (see figure 2K-M). Here, the data of figure 2K-M has been normalized only to the top three most stable housekeeping genes (Y45F10D.4, *pmp-3* and *lap-2*), and is highly similar to data normalized with the full set of five recommended housekeeping genes. See materials and methods for details on determination of stable genes using GeNorm. RM Two-way ANOVA with Geisser-Greenhouse and Šídák's multiple comparison test. For each ploidy, the mRNA levels were normalized such that levels without HS=1. The p-values for each factor of the Two-way ANOVA are indicated on the graph. Colours indicate matching independent biological replicates. Error bars = SEM.

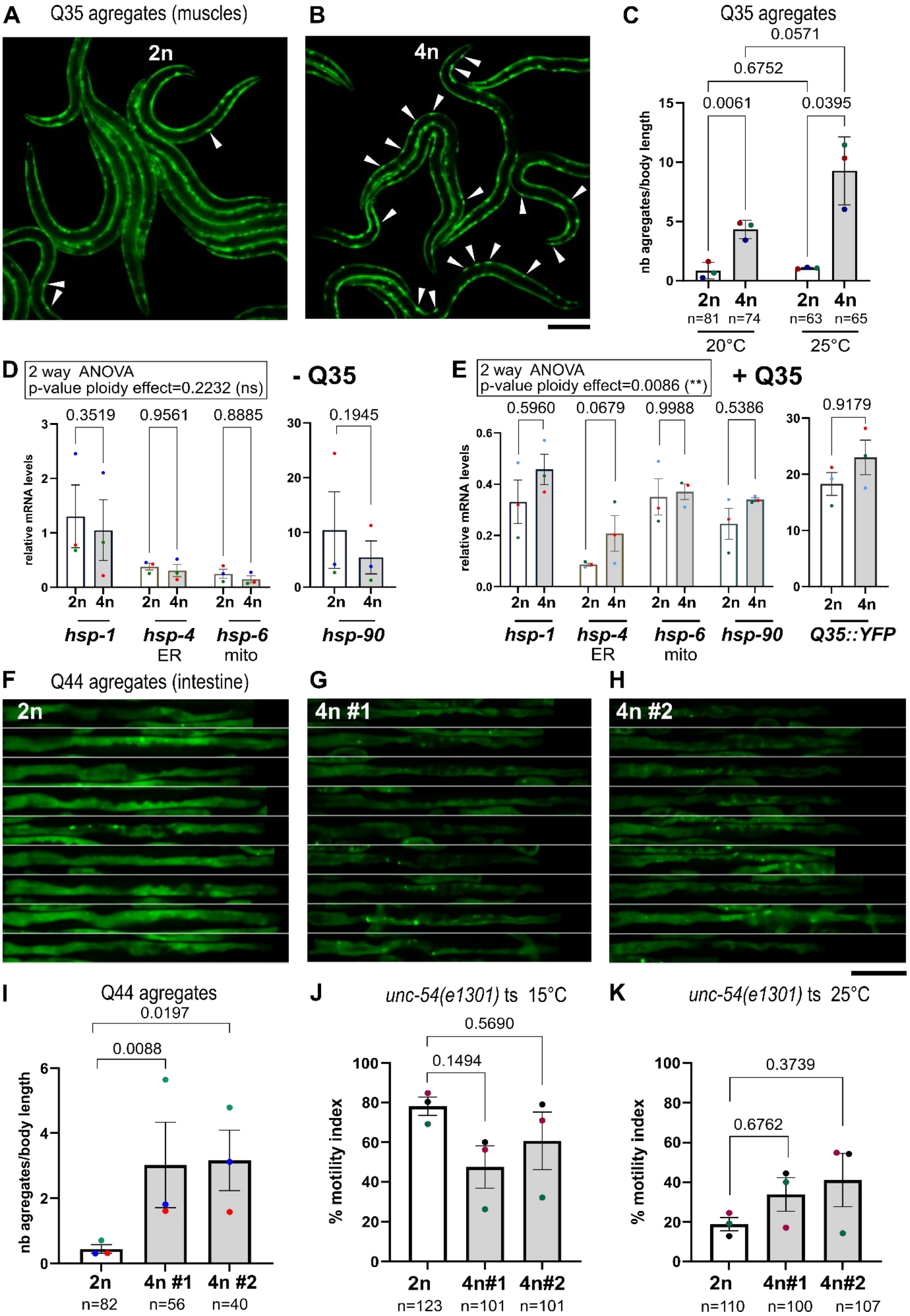


**Figure S2- Contrasting results obtained using different proteostasis sensors suggest a change in the behaviour of multicopy transgenes in neotetraploid animals.** (**A-C**) Tetraploids exhibit increased number of polyglutamine aggregates than diploids in muscles at both 20°C and 25°C. (**A-B**) Images of diploid AM140 (2n) and derived tetraploids MCL6 (4n), carrying *unc-54p*::Q35::YFP multicopy transgene expressed in body wall muscles, at day 2 of adulthood. Arrowheads indicate Q35 aggregates. Images were taken at Objective 10X on an epifluorescence microscope. Scale bar = 200 µm. (**C**) Quantification of Q35 aggregates in muscles of AM140 (2n) and derived MCL6 (4n) at day 2 of adulthood at both 20°C and 25°C. For each animal, the number of aggregates was normalised to the body length. Two-way ANOVA with Geisser-Greenhouse correction and uncorrected Fisher's LSD. Ploidy effect: p-value=0.025(*), temperature: p-value=0.0413(*), interaction: p-value=0.0881(ns). Adjusted pairwise p-values (Šídák's multiple comparisons test) are indicated on the graph. (**D-E**) Relative mRNA expression levels for constitutive molecular chaperones from the hsp-70 family: *hsp-1*, *hsp-4* (ER), *hsp-6* (mitochondria) and *hsp-90* measured by quantitative real-time PCR (qRT-PCR) in WT N2 (2n) and derived MCL2 (4n #2) animals at L4 stage in (**D**). In (**E**), the levels of those molecular chaperones were measured together with *yfp* mRNA levels in AM140 (2n) and derived MCL6 (4n) animals carrying the unc-54p::Q35::YFP mutlicopy transgene at the L4 stage. The geNorm algorithm[72] was used to determine the most stable set of housekeeping genes and the optimal number of housekeeping genes for comparisons at the L4 stage only: Y45F10D.4, *act-1*. Two-way ANOVA analysis was performed (Šídák's multiple comparison test). Comparisons between ploidies are indicated in Figure S2 and show no significant differences between diploids and tetraploids for comparisons for each target gene. However, considering all comparisons across target genes, two-way analysis comparing ploidies showed that mRNA was globally increased in tetraploids considering all target genes (p-value ploidy effect=0.0086, ns) in the presence of Q35::YFP transgene (E), but not in its absence (p-value ploidy effect=0.2232, ns), (D). (**F-I**) Tetraploids exhibit an increased number of polyglutamine aggregates than diploids in the intestine. (**F-H**) Images of intestines of diploid AM738 (2n) and derived tetraploids MCL20 (4n #1) and MCL21 (4n #2), carrying *vha-6p*::Q44::YFP multicopy transgene expressed in the intestine, at day 4 of adulthood. Images were taken at objective 10X on an epifluorescence microscope. Scale bar = 200 µm. (**I**) Quantification of Q44 aggregates in the intestine of AM738 (2n) and derived tetraploids MCL20 (4n #1) and MCL21 (4n #2) at day 4 of adulthood. For each animal, the number of aggregates was normalised to the body length. RM one-way ANOVA on log-transformed data: p-value=0.0028(**). Adjusted pairwise p-values (Tukey's multiple comparisons test) are indicated on the graph. N values indicate the number of animals assayed. (**J-K**) Motility index of temperature sensitive *unc-54(e1301)* proteostasis sensor in muscles. Diploid CB1301 (2n) and derived tetraploids MCL25 (4n #1) and MCL26 (4n #2) were assayed at both 15°C permissive temperature (J) or at 25°C restrictive temperature (K). Two-way ANOVA: p-value interaction ploidy x temperature= 0.0276(*), p-value temperature=0.0013(*), p-value ploidy=0.7110(ns). Pairwise adjusted p-values (Šídák's multiple comparison test) are indicated on the graph. In all panels, error bars indicate SEM, and n values indicate the number of animals assayed. Colours indicate matching independent biological replicates.

To investigate the consequences of autotetraploidy on proteostasis capacity, we generated tetraploid animals carrying the proteostasis sensor *unc-54p::*Q35::YFP, a multicopy transgene expressing stretches of polyglutamine (polyQ) expansion repeats fused to GFP in muscles. This transgene is used to monitor proteostasis capacity in muscles, by visualisation of polyglutamine aggregates when proteostasis capacity is compromised[73]. As shown in **Figure S2A-C**, neotetraploid animals carrying *unc-54p::*Q35::YFP exhibited a marked increase in Q35 aggregation at both temperatures assayed (20°C and 25°C) at day 2 of adulthood, indicative of reduced protein folding capacity in muscles. These data contrast with other results showing a modest increase in thermorecovery of tetraploids at both 20°C and 25°C (**Figure 2C**), suggesting a modest increase in proteostasis capacity of tetraploids at the whole animal level, when they do not carry any multicopy transgene. We investigated endogenous mRNA levels for several constitutive molecular chaperones in the presence or absence of the multicopy *unc-54p*::Q35:YFP transgene. While levels were globally unaffected by ploidy in the absence of the Q35::YFP transgene (p-value ploidy effect=0.2232), molecular chaperone levels were globally higher in animals carrying Q35::YFP (p-value ploidy effect=0.0086) with the highest effect observed on the endoplasmic reticulum (ER) chaperone *hsp-4* (p-value=0.0679), a sign of ER stress. Q35::YFP is a highly expressed mutlicopy transgene (see difference in scale in **Figure S2E**), and its levels seemed higher in tetraploids (even if not significantly so). It is possible that higher levels of this transgene in tetraploids are negatively affecting cellular protein folding capacity. To determine if proteostasis capacity was also affected by the expression of multicopy transgenic proteostasis sensors in other tissues, we tetraploidized animals carrying *vha-6p*::Q44::YFP, expressed in the intestine. At day 4 of adulthood neotetraploids also exhibited a significant increase in intestinal Q44 aggregation at 20°C in two independent lines (**Figure S2F-I**). To disentangle the effects of multicopy transgenic proteostasis sensors on protein folding capacity in neotetraploids, we assayed proteostasis capacity in muscles in the absence of any multicopy transgene, using the *unc-54(e1301)* temperature sensitive (ts) mutation in heavy-chain myosin with phenotypic consequences in muscles. In *unc-54* ts mutants, a decrease in protein folding capacity is correlated with motility defects. Our results indicate that tetraploidy does not affect the motility of *unc-54(e1301)* mutants (p-value ploidy effect=0.7110), but there is an interaction between temperature and ploidy (p-value interaction=0.0276) with a slight decrease in motility at 15°C (**Figure S2J**) and a small improvement at 25°C (**Figure S2K**).

These results suggest that tetraploidy affects multicopy transgenes, possibly by affecting their regulation and leading to a decrease in protein folding capacity. To test this hypothesis, we generated neo-tetraploid animals carrying the heat-inducible transcriptional reporter of the constitutive and heat inducible molecular chaperon *hsp-*$90_p$::GFP, either as a multicopy or single copy transgene. The fluorescence levels of *hsp-*$90_p$::GFP were indeed increased in tetraploids carrying the multicopy transgene (**Figure S3A,B,E**), but not in the case of the single-copy transgene (**Figure S3C,D, F**).

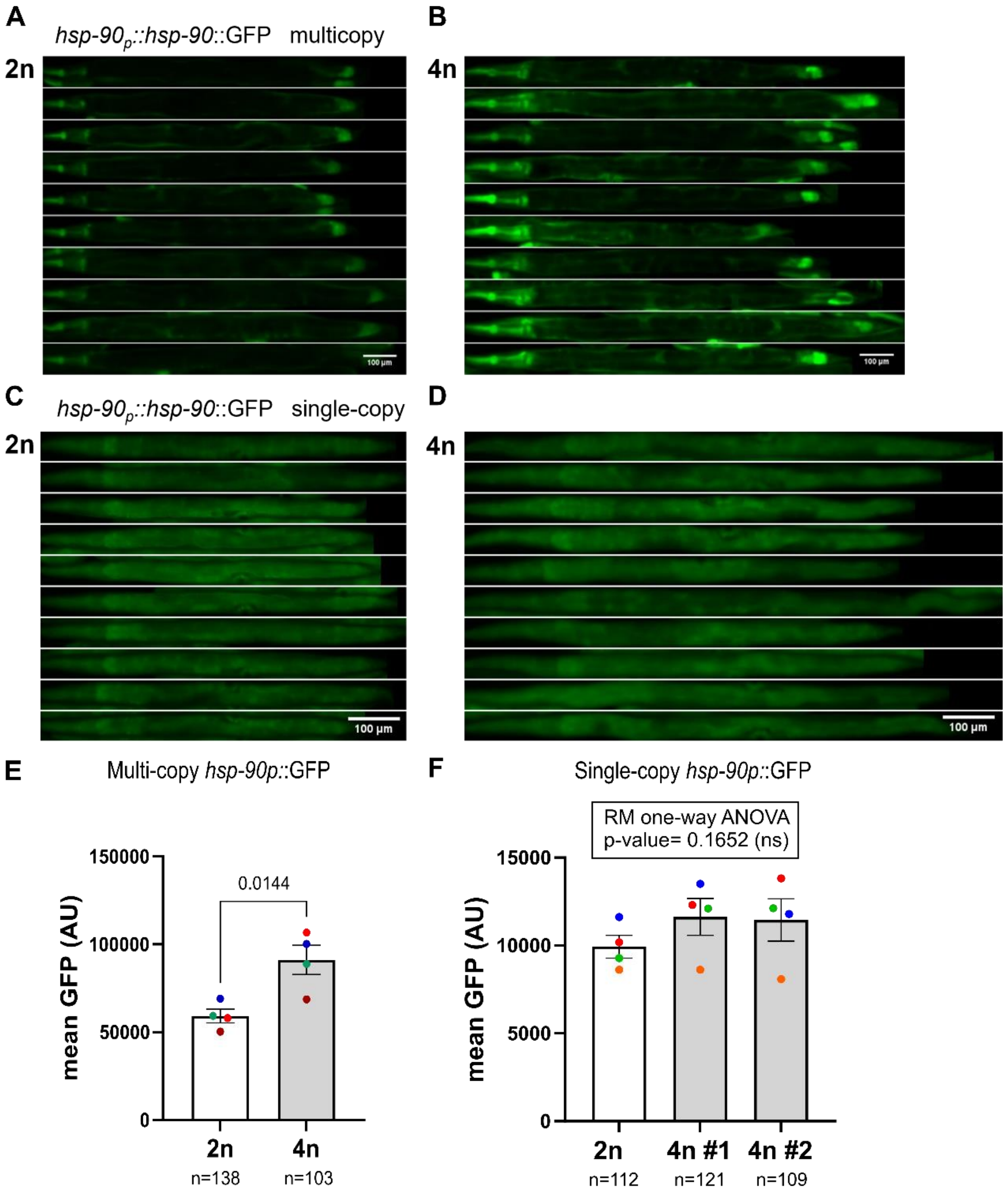


**Figure S3- Tetraploidy differentially affects multicopy and single-copy transgenes.** The fluorescence levels of multicopy transcriptional *hsp-90p::*GFP reporter are increased in tetraploids, whereas fluorescence levels of single-copy *hsp-90::*GFP reporter are comparable in diploids and tetraploids animals. **(A-B)** Micrographs of multicopy *hsp-90p::*GFP reporter in diploid BCN1049 (A) and tetraploid MCL11 (B) animals. **(C-D)** Micrographs of single-copy *hsp-90p::*GFP reporter in diploid BCN1082 (C) and tetraploid MCL36 (4n#1) (D) animals. Scale bar: 100 µm. **(E)** Mean fluorescence intensity levels (arbitrary units) for multicopy *hsp-90p::*GFP reporter in diploid (BCN1059) and tetraploid (MCL1) animals. Paired t-test. **(F)** Mean fluorescence intensity levels (arbitrary units) for single-copy *hsp-90p::*GFP reporter in diploid (BCN1082) and tetraploid (MCL38 4n#1 and MCL36 4n#2) animals. RM One-way ANOVA with Geisser-Greenhouse correction and Tukey's multiple comparison test. The p-value is indicated above the graph. Colours indicate matching independent biological replicates. The number of animals quantified in (E) and (F) is indicated below the x-axis.

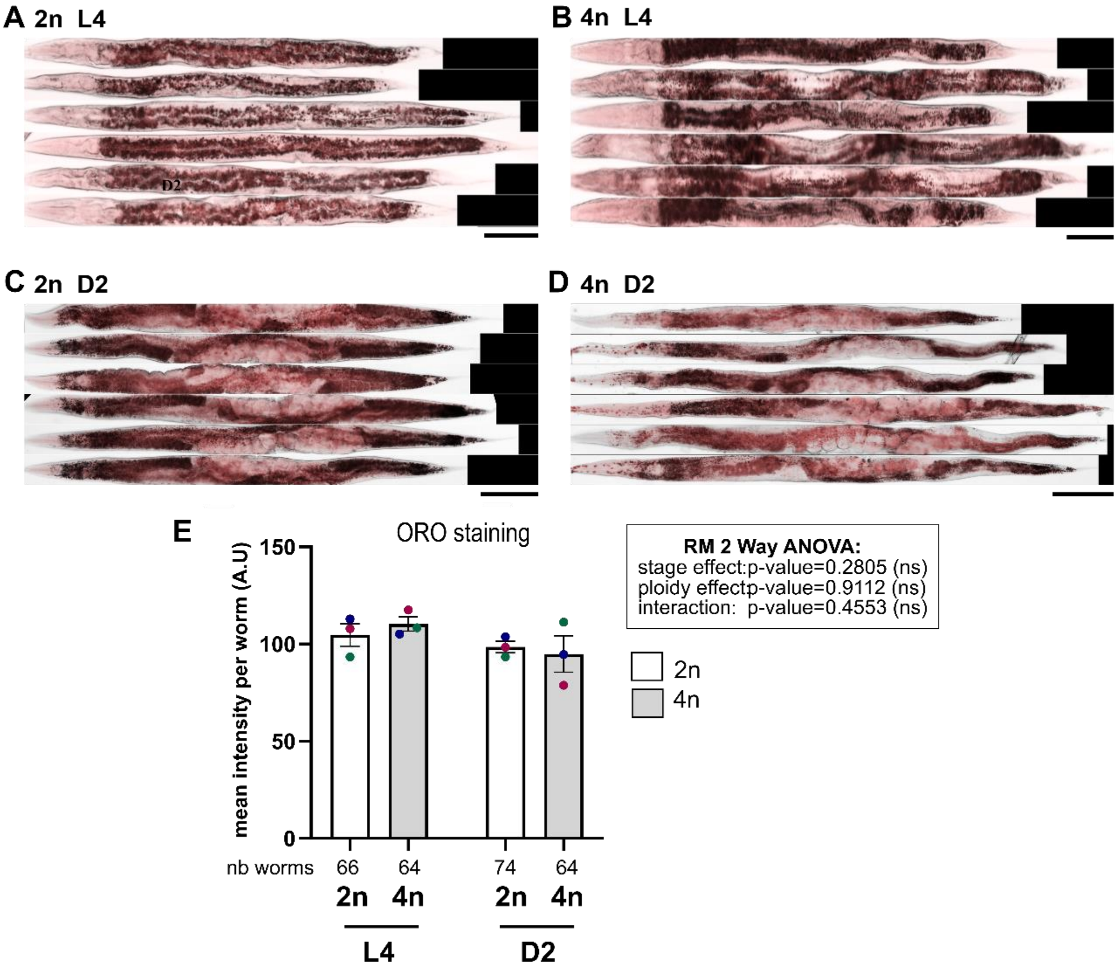


**Figure S4- ORO staining shows similar levels of neutral lipids in diploids and tetraploids under regular conditions. (A-D)** Micrographs L4 larvae diploid N2 (A), tetraploid MCL2 (B), and day 2 adults diploid N2 (C) and tetraploid MCL2 animals stained with neutral lipid dye Oil-Red-O (ORO), in the absence of cold shock. Scale bar: 100 µm. **(E)** Mean ORO intensity levels of diploids and tetraploids at either L4 or day 2 adult stage. RM Two-way ANOVA with Geisser-Greenhouse correction and Šídák's multiple comparison test. The p-values of the main effects are indicated on the graph. Colours indicate matching independent biological replicates. The number of animals quantified in (E) and (F) is indicated below the x-axis.

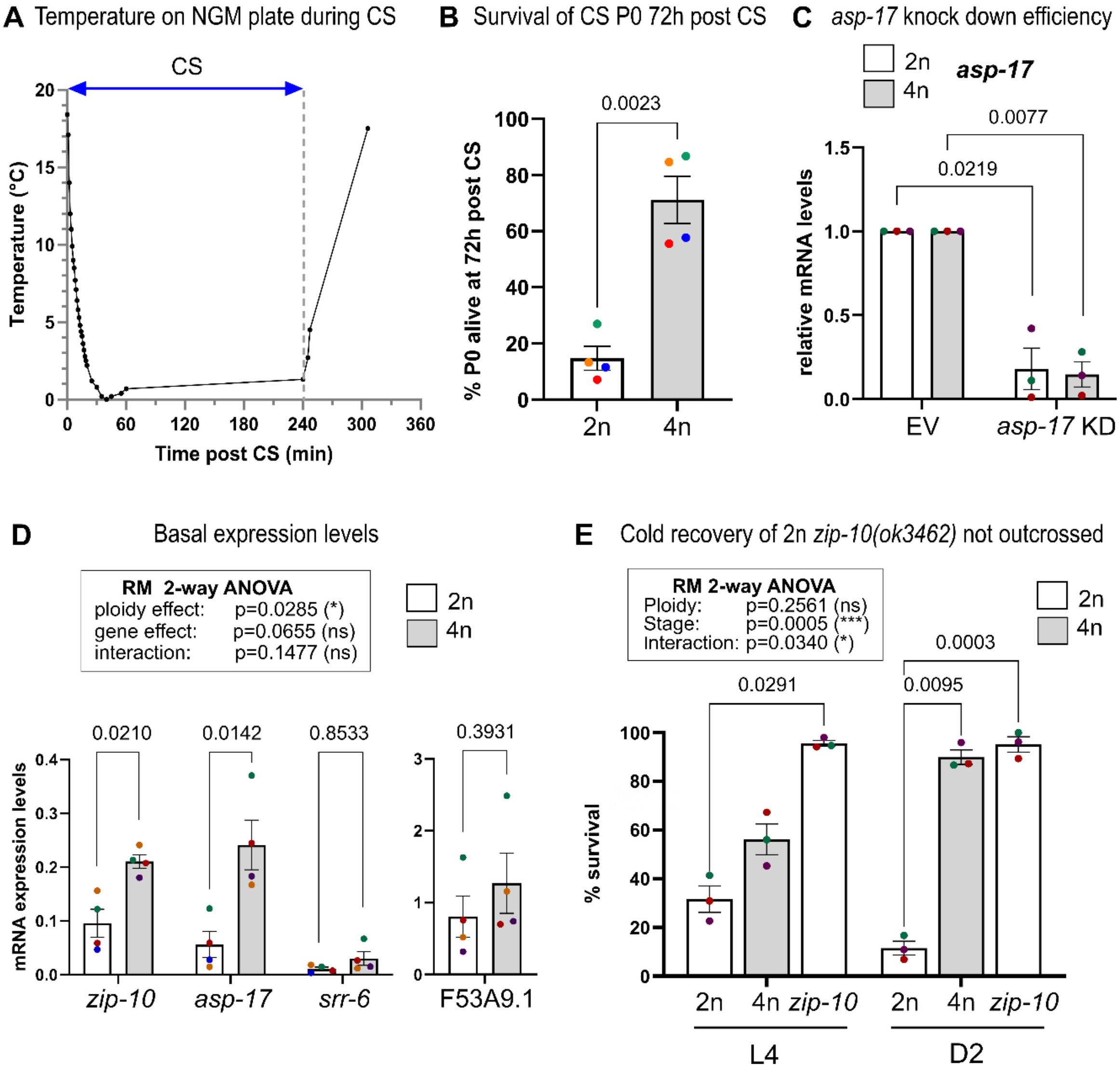


**Figure S5-Data associated with Figure 3 and Figure 6. Temperature during CS. Expression levels of cold-recovery induced mRNAs are upregulated under basal conditions in tetraploids. RNAi knock-down efficiency for *asp-17.* Survival upon CR in *zip-10(ok3462)* non-**outcrossed **mutant. (A)** Temperature on the surface of a 6cm NGM plate during and after a four-hour cold shock in an icebox. The temperature drops to 1°C within 30 minutes and stay between 0-1°C for the remaining of the cold shock. **(B)** Survival of cold shocked P0 of diploid N2 and derived tetraploid MCL2 at 72h post cold shock. Paired t-test. **(C)** Relative expression levels of the transcription factor *zip-10*, its target *asp-17*, and cold-induced *srr-6* and F53A5.9, in the absence of CS in diploid N2 and tetraploid MCL2 day 2 adults. For clarity (different scales), F53A9.1 was represented on a separated graph. RM Two-way ANOVA with Geisser-Greenhouse correction and Šídák's multiple comparisons test. In (B) and (C) levels of mRNA in day 2 adults were normalized to the housekeeping genes (*klp12, lap-2, act-1* and *pmp-3*). **(D)** Efficiency of *asp-17* RNAi knock-down in diploid and tetraploid animals. Upon *asp-17* RNAi, asp-17 mRNA levels are decreased by 82% in N2 diploids and by 85% in MCL2 tetraploids. Levels were normalized so that levels in control (Empty Vector) =1. One sample t-test. **(E)** Survival (% alive) upon cold recovery of diploid N2 (2n), derived tetraploid MCL2 (4m #2) and diploid not outcrossed *zip-10(ok3462)* loss of function mutant at L4 or Day 2 adult stage (D2). RM Two-way ANOVA with Geisser-Greenhouse correction. Only significant pairwise adjusted p-values (Šídák's multiple comparisons test) are indicated on the graph. Colours indicate matching independent biological replicates.

**Table S1- Primers used in this study.**

| **qRT-PCR primers** | | | |
|---|---|---|---|
| **mRNA target** | **Identifier For/Rev** | **Forward** | **Reverse** |
| *act-1* | act-1 F/R | GCTGGACGTGATCTTACTGATTACC | GTAGCAGAGCTTCTCCTTGATGTC |
| *ama-1* | ama-1 F/R | TGATCCGATGAATGATGGAA | TTCCATTCTGCGTTGATGTC |
| *asp-17* | asp-17 F/R | TGGGGTCACTTATGTTCCGC | CCGTGTCGGAAATTACCTGATTG |
| *asp-17 RNAi KD* | asp-17 KD F/R | TCGGTAACTTCACTGTGGGG | TCCAGTGTCAAGGACCAGG |
| *cdc-42* | cdc-42 F/R | TCCACAGACCGACGTGTTTC | AGGCACCCATTTTTCTCGGA |
| *F53A9.5* | F53A9.5 F/R | ACTACGGAAACGGAGGATAC | TGGCCGTGATGATGATGATG |
| *GFP* | GFP F/R | TGTTCCATGGCCAACACTTG | CCTGTACATAACCTTCGGGCA |
| *gpd-2* | gpd-2 F/R | CGGAGTCTTCACCACCATCG | CGACGAACATTGGAGCATCA |
| *hsp-1* | hsp-1 F/R | CACTGTTTTCGATGCCAAACG | TCCTTCGGCAGAGATGACCT |
| *hsp-16.1/11* | hsp-16.1 F/R | ATGGCTCAGATGGAACGTCA | TGGCTTGAACTGCGAGACAT |
| *hsp-16.2* | hsp-16.2 F/R | TCCATCTGAGTCTTCTGAGATTGTT | TGATAGCGTACGACCATCCAAA |
| *hsp-4* | hsp-4 F/R | TTCAACAAGACATCAAGCACTGG | GGCAGAGACTTCTTCAGGAGTGA |
| *hsp-6* | hsp-6 F/R | GATTGGATAAGGACGCTGGAGA | CCGTTGGTGGACTTGACCTC |
| *hsp-70 (C12C8.1)* | hsp-70 F/R | TCGATGAAGTTGTCTTGGTTGG | AGGCTACTGCTTCGTCTGGATT |
| *hsp-90* | hsp-90 F/R | ATTCGCTACCAGGCACTCAC | TGGTAAGGGTCTTTTCCTCCT |
| *ife-1* | ife-1 F/R | CAGCGTCTGGACTAAGGATTGC | GGATCACATCGAACAGTGGCTT |
| *ire-1* | ire-1 F/R | TACTTGCCACCACGGAGACC | CGTTGCCATCGTCATCATTG |
| *klp-12* | klp-12 F/R | TGCGTAACAATAATCGAAAGCA | CTGACAAAACACCCCTTGCG |
| *lap-2* | lap-2 F/R | GCCGTGACAAGTACGGATCT | CACATGTACCCGACAGCCTT |
| *srr-6* | srr-6 F/R | ATCCGCGTTGTTAGAGAAACC | TGAACTGCTGAATCCACTGGC |
| *pmp-3* | pmp-3 F/R | GTTCCCGTGTTCATCACTCAT | ACACCGTCGAGAAGCTGTAGA |
| *tsn-1* | tsn-1 F/R | GTCAAGGAAAAGTTGCTGATGAA | GTTCCATTTTCCGCGTCCTG |
| *unc-16* | unc-16 F/R | GGAGTATATCGACCCAGACATGATT | AGAGTTCAACAGATTGGTCTTCG |
| *vit-2* | vit-2 F/R | TCCATCAAGAGCCACATCAAGA | CGAACTCAGCCTTGTCTCCA |
| *vit-5* | vit-5 F/R | AGAATCTGAGGTTCATTCCGTT | GTCCAGAAACCTTCTTGATCTCTCT |
| *vit-6* | vit-6 F/R | ATATTCAAGAATCCTCATTCCGCGC | CGTGAGATTCCTTGTGGGGT |
| *YFP* | YFP F/R | CTACCCCGACCACATGAAGC | CTTGTAGTTGCCGTCGTCCT |
| *Y45F10D.4* | Y45F10D.4 F/R | GTCGCTTCAAATCAGTTCAGC | GTTCTTGTCAAGTGATCCGACA |
| *zip-10* | zip-10 F?R | ATCCAGCTCGAGATGCTCTTC | ATAAAGACGGCGATGACGCT |
| **genotyping primers** | | | |
| **mRNA target** | **Identifier** | **Forward** | **Reverse** |
| zip-10 | ok3462 flank. F/R | GCACAACTCGGGTGCTCATA | AAGAAACGAGGTGGGGATGG |
| zip-10 | ok3462 withinF/R | TCAATCTGCCTGTTTTGCCA | CACGGTACTGGCGAGCATAA |

**Table S2- Survival of diploids and neotetraploids exposed to *P. aeruginosa*.**

| Strain | Ploidy | FUdR | Rep. | Nb. deaths | Nb. censored | median survival | P-value (vs 2n) Log-rank (Mantel-Cox) | Hazard ratio (2n/4n |
|---|---|---|---|---|---|---|---|---|
| N2 | 2n | yes | 1 | 99 | 75 | 4 | | |
| MCL2 | 4n | yes | 1 | 98 | 46 | 4 | 0.002 (**) | 0.7261 |
| N2 | 2n | yes | 2 | 81 | 41 | 4 | | |
| MCL2 | 4n | yes | 2 | 103 | 11 | 4 | 0.7546 (ns) | 1.031 |
| EJ1171 | 2n | no | 1 | 103 | 59 | 6 | | |
| MCL22 | 4n | no | 1 | 83 | 82 | 6 | 0.5862 (ns) | 0.9358 |
| EJ1171 | 2n | no | 2 | 109 | 46 | 7 | | |
| MCL22 | 4n | no | 2 | 86 | 72 | 7 | 0.9263 (ns) | 0.9838 |

**Table S3- Lifespan of diploids and neotetraploids raised at 20°C or 25°C.**

| Strain | Ploidy | Temp. | Rep. | Nb. deaths | Nb. censored | median survival | P-value (vs 2n) Log-rank (Mantel-Cox) | Hazard ratio 2n/4n |
|---|---|---|---|---|---|---|---|---|
| N2 | 2n | 20°C | 1 | 47 | 103 | 21 | | |
| MCL1 (4n#1) | 4n | 20°C | 1 | 75 | 73 | 17 | <0.0001(****) | 0.5483 |
| MCL2 (4n#2) | 4n | 20°C | 1 | 63 | 87 | 17 | <0.0001(****) | 0.4508 |
| N2 | 2n | 20°C | 2 | 111 | 49 | 21 | | |
| MCL1 (4n#1) | 4n | 20°C | 2 | 74 | 76 | 16 | <0.0001(****) | 0.52 |
| MCL2 (4n#2) | 4n | 20°C | 2 | 94 | 79 | 16 | <0.0001(****) | 0.5504 |
| N2 | 2n | 20°C | 3 | 76 | 68 | 22 | | |
| MCL1 (4n#1) | 4n | 20°C | 3 | 79 | 87 | 17 | <0.0001(****) | 0.526 |
| MCL2 (4n#2) | 4n | 20°C | 3 | 79 | 70 | 15 | <0.0001(****) | 0.4619 |
| N2 | 2n | 25°C | 1 | 106 | 43 | 10 | | |
| MCL2 (4n#2) | 4n | 25°C | 1 | 84 | 81 | 12 | 0.0254(*) | 1.315 |
| N2 | 2n | 25°C | 2 | 77 | 74 | 12 | | |
| MCL2 (4n#2) | 4n | 25°C | 2 | 63 | 54 | 15 | 0.2775(ns) | 1.161 |
| N2 | 2n | 25°C | 3 | 49 | 95 | 9 | | |
| MCL1 (4n#1) | 4n | 25°C | 3 | 56 | 93 | 11 | 0.1095(ns) | 1.299 |
| N2 | 2n | 25°C | 4 | 72 | 85 | 11 | | |
| MCL1 (4n#1) | 4n | 25°C | 4 | 93 | 65 | 11 | 0.8008(ns) | 1.033 |
| MCL2 (4n#2) | 4n | 25°C | 4 | 80 | 86 | 13 | 0.0869(ns) | 1.248 |
| N2 | 2n | 25°C | 5 | 79 | 36 | 9 | | |
| MCL1 (4n#1) | 4n | 25°C | 5 | 94 | 44 | 9 | 1.1707(ns) | 0.845 |
| MCL2 (4n#2) | 4n | 25°C | 5 | 95 | 38 | 9 | 1.0102(*) | 0.7009 |
